# GRouNdGAN: GRN-guided simulation of single-cell RNA-seq data using causal generative adversarial networks

**DOI:** 10.1101/2023.07.25.550225

**Authors:** Yazdan Zinati, Abdulrahman Takiddeen, Amin Emad

**Affiliations:** Department of Electrical and Computer Engineering, McGill University, Montreal, QC, Canada; Mila, Quebec AI Institute, Montreal, QC, Canada; The Rosalind and Morris Goodman Cancer Institute, Montreal, QC, Canada

## Abstract

We introduce GRouNdGAN, a gene regulatory network (GRN)-guided causal implicit generative model for simulating single-cell RNA-seq data, *in-silico* perturbation experiments, and benchmarking GRN inference methods. Through the imposition of a user-defined GRN in its architecture, GRouNdGAN simulates steady-state and transient-state single-cell datasets where genes are causally expressed under the control of their regulating transcription factors (TFs). Training on three experimental datasets, we show that our model captures non-linear TF-gene dependences and preserves gene identities, cell trajectories, pseudo-time ordering, and technical and biological noise, with no user manipulation and only implicit parameterization. Despite imposing rigid causality constraints, it outperforms state-of-the-art simulators in generating realistic cells. GRouNdGAN learns meaningful causal regulatory dynamics, allowing sampling from both observational and interventional distributions. This enables it to synthesize cells under conditions that do not occur in the dataset at inference time, allowing to perform *in-silico* TF knockout experiments. Our results show that *in-silico* knockout of cell type-specific TFs significantly reduces cells of that type being generated. Interactions imposed through the GRN are emphasized in the simulated datasets, resulting in GRN inference algorithms assigning them much higher scores than interactions not imposed but of equal importance in the experimental training dataset. Benchmarking various GRN inference algorithms reveals that GRouNdGAN effectively bridges the existing gap between simulated and biological data benchmarks of GRN inference algorithms, providing gold standard ground truth GRNs and realistic cells corresponding to the biological system of interest. Our results show that GRouNdGAN is a stable, realistic, and effective simulator with various applications in single-cell RNA-seq analysis.

## Introduction

Unraveling gene regulatory interactions, often represented as a gene regulatory network (GRN), plays a crucial role in studying biological processes under different conditions^1–3^, simulating knockdown and knockout experiments^4, 5^, and identifying therapeutic drug targets^6, 7^. Many algorithms have been proposed for GRN reconstruction using bulk or single-cell RNA sequencing data (scRNA-seq), alone^8–14^ or with other modalities^15–18^. While these advances have provided great biological insights, evaluating the performance of GRN Inference algorithms remains challenging^19, 20^ due to the lack of reliable ground truth for the biological processes under study. Existing evaluation approaches often resort to curated databases^21–23^. However, the regulatory interactions in these databases are aggregated from a wide range of datasets and are not specific to a biological system, making them not ideal benchmarks due to context specificity of gene regulation. Another strategy is to verify regulatory interactions by conducting perturbation experiments on the system under study^20^. However, this approach is tedious, lengthy, and expensive. Another approach is to employ scRNA-seq simulators. Despite great progress in this domain, most simulators lack the essential properties for this task, such as the preservation of gene identities and simulation based on a user-provided ground truth GRN. For example, scGAN^24^, cscGAN^24^, scDESIGN2^25^, and SPARSIM^26^ capture inter-gene correlations in their datasets. However, since they do not explicitly impose a known GRN (that can act as the ground truth), they are not suitable for benchmarking GRN inference methods.

A small subset of simulators (e.g., BoolODE^19^, SERGIO^4^, and GeneNetWeaver (GNW)^27^) explicitly incorporate GRNs capturing transcription factor (TF)-gene dynamics. GNW was used to generate benchmarks for DREAM challenges^27–29^. However, being a bulk RNA-seq tool, its simulated datasets do not replicate the distribution of experimental scRNA-seq datasets nor exhibit their statistical properties, despite attempts to adapt it for this purpose by externally inducing dropout events^10, 30^. SERGIO and BoolODE were designed to simulate scRNA-seq data using stochastic differential equations (SDEs)^4, 19^ and have been used to benchmark a variety of GRN inference methods. However, often a mismatch between the benchmarking results based on experimental and their simulated data have been reported, which may be attributed to differences between the simulated and experimental datasets. For example, in the BEELINE study (using BoolODE), some of the top ranking methods on simulated datasets reported near random performance on experimental datasets^19^. Both simulators enable the user to simulate more realistic datasets by carefully selecting the values of the SDE parameters. Moreover, using a reference dataset, SERGIO allows fine tuning the added technical noise in an iterative procedure until the generated and reference datasets are matched^4^. While these steps can help improve the resemblance to real datasets, they are often suboptimal and pose an undue burden on the user: for example, in SERGIO, the user must evaluate the similarity based on five different statistics and change three parameters iteratively until desired resemblance is achieved (which itself can be subjective). In addition, since the GRN is imposed on the “clean” dataset, but technical noise is added afterwards, this step may change the encoded causal relationships in a non-trivial way (which may explain why the performance of GRN inference methods dropped close to random when applied to the noisy dataset^4^). Furthermore, SERGIO and BoolODE do not readily preserve gene identities and make simplifying assumptions regarding cooperative regulation (see Discussion). These shortcomings demonstrate the need for simulators capable of generating realistic scRNA-seq data that retain the regulatory dynamics specified by a user-defined GRN. Importantly, with the growing interest in causal inference, simulators that impose *causal* GRNs are of great need.

GRouNdGAN (<u>GR</u>N-guided in silic<u>o</u> sim<u>u</u>lation of single-cell R<u>N</u>A-seq <u>d</u>ata using Causal <u>g</u>enerative <u>a</u>dversarial <u>n</u>etworks) is a causal implicit generative model for GRN-guided simulation of scRNA-seq data inspired by CausalGAN^31^. Given an input GRN and a reference dataset, it can be trained to generate simulated data that is both indistinguishable from the reference data and faithful to the causal regulatory interactions of the input GRN. Unlike model-based simulators that rely on simplifying assumptions regarding co-regulatory patterns, in GRouNdGAN these patterns are *learned* through complex functions. This allows it not to compromise on the underlying complexity of the system and to model elaborate regulatory dynamics. GRouNdGAN provides state-of-the-art performance in realistic scRNA-seq data generation, while preserving gene identities, user-defined causal gene regulatory interactions, and cellular dynamics (e.g., lineage trajectory and pseudo-time ordering). This is achieved through implicit parameterization without the need for manual fine-tuning. Using GRouNdGAN, we benchmark seven GRN inference methods and find the results to be aligned with BEELINE’s experimental results^19^. Furthermore, the causal structure of GRouNdGAN enables it to be used for sampling from both interventional and observational data distributions, enabling *in silico* knockout experiments.

## Results

### GRouNdGAN generates scRNA-seq data using causal generative adversarial networks

GRouNdGAN is a deep learning model that generates scRNA-seq data while imposing a user-defined causal GRN to describe the regulatory relationships of the genes and TFs. Its architecture builds on the causal generative adversarial network^31^ and includes a causal controller, target generators, a critic, a labeler and an anti-labeler (Figure 1). Training includes two steps. First, the causal controller that generates TF expression values is pre-trained (Figure 1A) as the generator of a Wasserstein GAN (WGAN)^32^ (Methods). In the second step (Figure 1B), TF expression values (generated by the pre-trained causal controller) and randomly generated noise are provided as input to the target generators that produce target genes’ expressions, while incorporating the TF-gene relationships of the input GRN. To achieve this, as input, each target generator only accepts a noise value and the generated expression values of TFs that causally regulate it (Figure 1B, Methods). The generated expression of genes and TFs are then fed to a library-size normalization (LSN) layer^24^.

**Figure 1:**
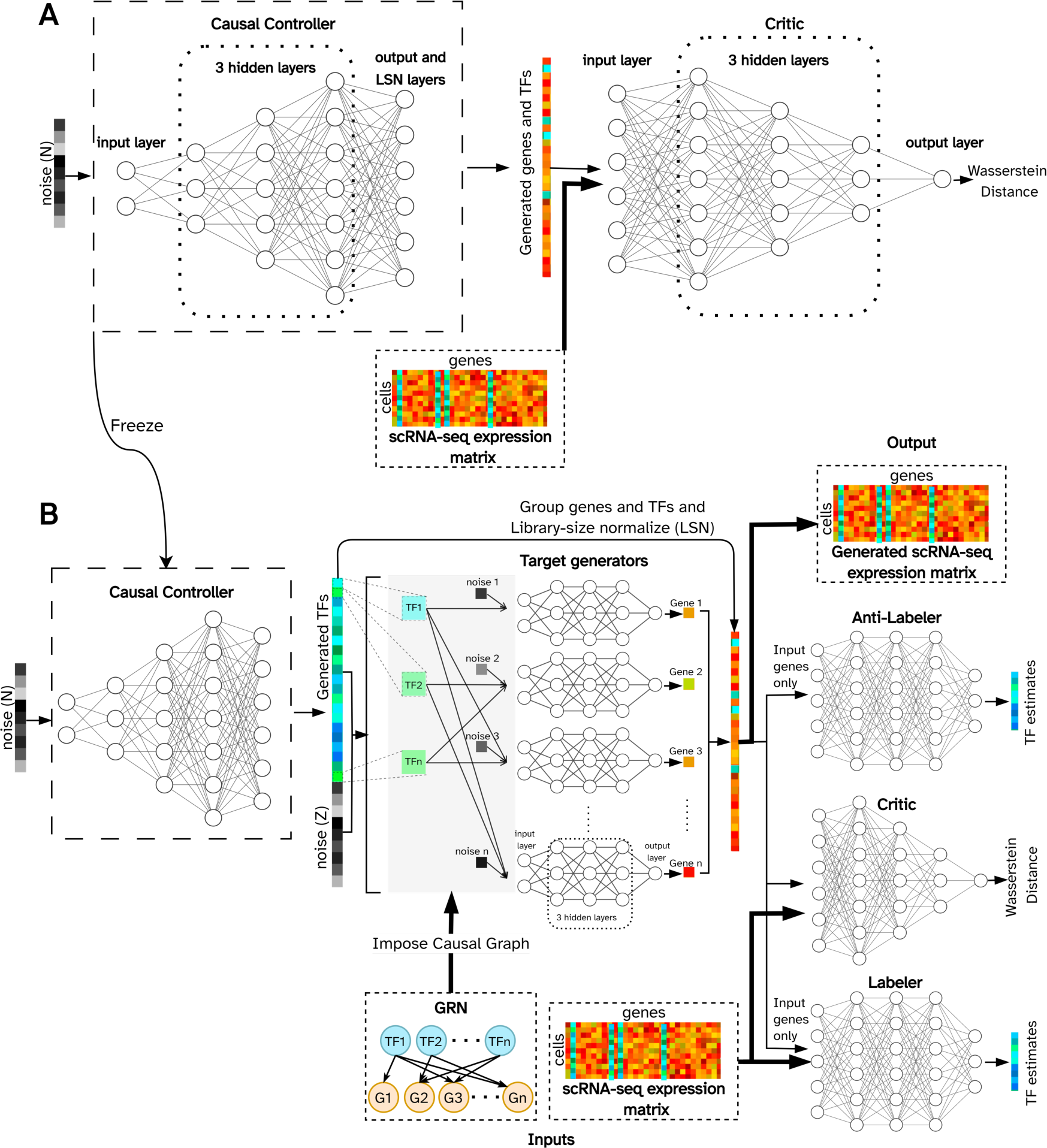
Architecture and training procedure of GRouNdGAN. A) A WGAN is pre-trained to generate realistic simulated cells. B) The LSN layer of the generator of the trained WGAN (panel A) is removed, its weights are frozen, and is used as the causal controller to generate unnormalized TF expression values. These TF expression values along with a noise vector are provided as input to the target generators, following the provided causal GRN. The generated gene and TF expression values are reorganized and passed through the LSN layer. The normalized simulated expression vectors and experimental reference data are then passed to the critic to estimate Wasserstein distance between the reference and the generated data distributions. The anti-labeler estimates TF values based on generated target gene expressions. The labeler performs a similar task, but in addition to receiving generated values, it also utilizes target gene expression values from the reference data. Labeler and anti-labeler ensure that the causal GRN is incorporated by the target generators. Details of the model are provided in Methods.

The critic’s role is to quantify the Wasserstein distance between the reference and simulated data. The target generators are trained in an adversarial manner to generate realistic datapoints indistinguishable from the reference datapoints by the critic. The labeler/anti-labeler estimate the causal controller’s TF expression values *only* from the target genes’ expressions to ensure that the generated causal TF-gene dependencies are encoded. Details are provided in Methods, the architectural choices in Tables S1-S2, and an ablation study in File S1 and Table S3.

### GRouNdGAN generates realistic scRNA-seq data

We trained GRouNdGAN on three datasets. The first dataset contained scRAN-seq profiles of 68579 human peripheral blood mononuclear cells (PBMCs) from 10x Genomics^33^ corresponding to eleven cell types (“PBMC-All”). We formed a dataset of the most common cell type in the PBMC-All dataset containing 20773 CD8+ Cytotoxic T-cells (“PBMC-CTL”). Additionally, we obtained the scRNA-seq (MARS-seq) profile of 2730 cells corresponding to differentiation of hematopoietic stem cells to different lineages from mouse bone marrow^34^ (“BoneMarrow”). We used GRNBoost2^8^ to identify fifteen TFs for each gene to form the input GRN. Note that these identified interactions may contain both spurious correlations and causal relationships in the reference dataset. However, when they are used as input to GRouNdGAN, they will correspond to causal interactions in the *simulated* dataset (see Kocaoglu et al. for theoretical details of causality in this architecture^31^). For each dataset, we trained GRouNdGAN on a randomly selected training set and evaluated on a held-out test set (Methods, Table S4). Figures 2A-2B and S1-S3 show the t-SNE plots of reference and simulated cells, qualitatively revealing their similarity.

**Figure 2:**
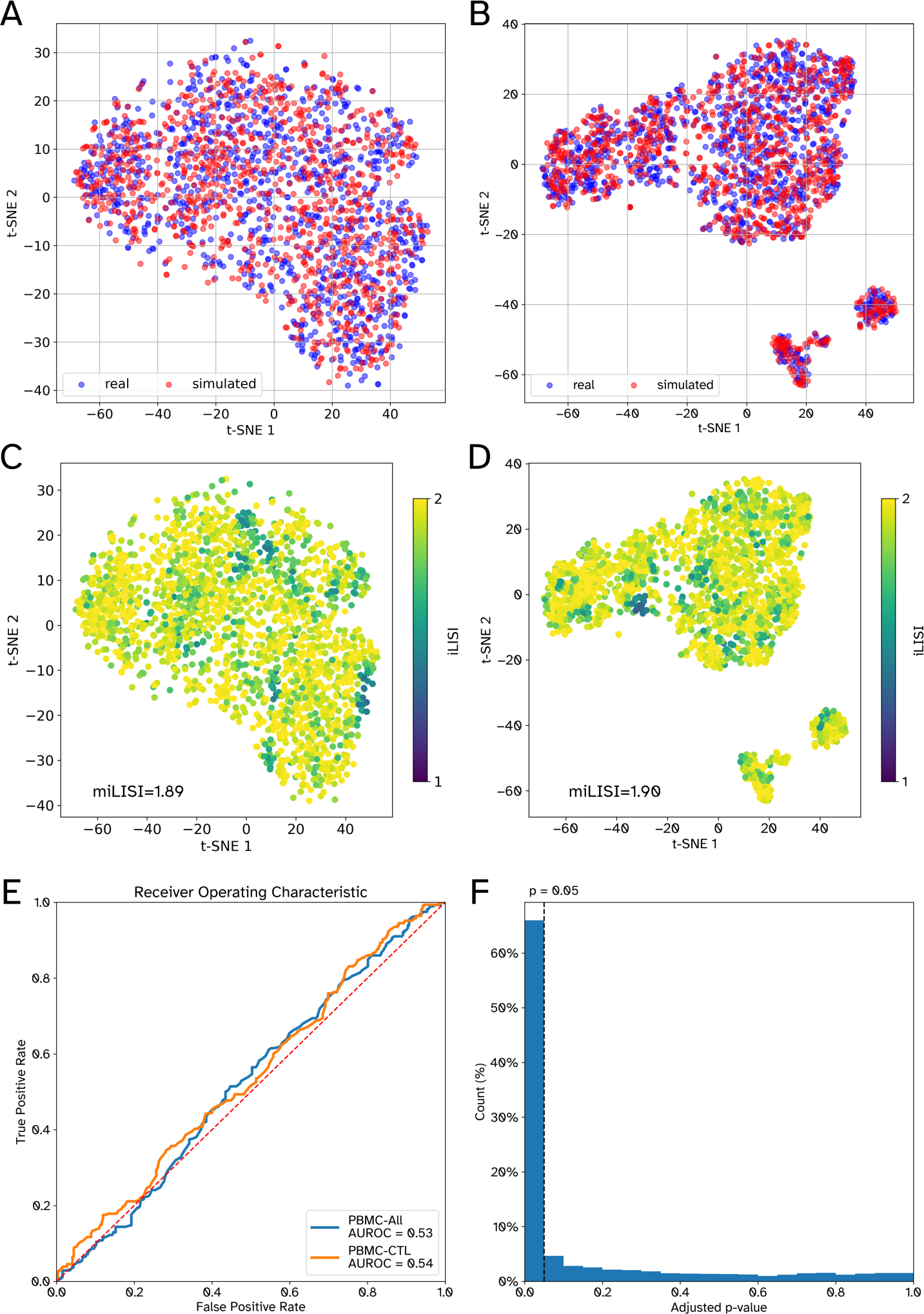
Performance of GRouNdGAN in generating realistic scRNA-seq data. All plots correspond to 1000 real (experimental) held-out test cells and 1000 simulated cells. Each gene in the imposed GRN of GRouNdGAN is regulated by 15 TFs (identified using GRNBoost2 from the experimental training set). Panels A and B depict the t-SNE plots of real (red) and simulated (blue) cells for PBMC-CTL and PBMC-All, respectively, while panels C and D also show their iLISI distributions. E) The ROC curve of a Random Forest classifier in distinguishing between simulated and real cells. F) The distribution of adjusted p-values corresponding to the TF perturbation study on the PBMC-CTL dataset. The p-values are obtained using two-sided Wilcoxon signed ranked tests and are adjusted for multiple hypotheses following the Benjamini-Hochberg procedure.

We quantitatively assessed their similarity using Euclidean distance, Cosine distance, maximum mean discrepancy (MMD)^35^, mean integration local inverse Simpson’s index (miLISI)^36^, and the area under the receiver operating characteristic curve (AUROC) of a random forests (RF) classifier distinguishing simulated and experimental cells (Methods). As a “control”, we calculated these metrics using two halves of the reference test set. Table 1 shows the performance of GRouNdGAN, control, and three state-of-the-art simulators for the PBMC-CTL dataset (see Table S4 for training set performance). Note that to assess data quality systematically and fairly, we only included methods that automatically generate scRNA-seq data resembling experimental data, since the quality of data generated by methods that rely on the user’s judgement for distribution matching (e.g., SERGIO and BoolODE) is user dependent and subjective.

**Table 1:** Performance of different simulators in generating realistic scRNA-seq data using the PBMC-CTL dataset. The metrics are calculated between a simulated dataset of 1000 cells and the held-out test set of 1000 real cells (see Table S4 for training set performance). Each gene in the imposed GRN of GRouNdGAN is regulated by 15 TFs (constructed using GRNBoost2 from the experimental training set). For the first three metrics, a value closer to zero is preferred, for RF AUROC a value closer to 0.5 is preferred, and for miLISI a value closer to 2 is preferred. Best performance values (excluding control) are in bold-face. The control shows the value of the metrics comparing the two halves of the real dataset. For the first two metrics, the values correspond to the distance of the mean centroids of the real and simulated cells.

| Simulator | Cosine distance | Euclidean distance | MMD | RF AUROC | miLISI |
| --- | --- | --- | --- | --- | --- |
| GRouNdGAN | **0.00057** | **182** | **0.026** | **0.54** | **1.891** |
| scGAN<sup>24</sup> | 0.00095 | 222 | 0.031 | 0.59 | 1.888 |
| scDESIGN2<sup>25</sup> | 0.00100 | 229 | 0.065 | 0.76 | 1.736 |
| SPARsim<sup>26</sup> | 0.00104 | 235 | 0.309 | 0.95 | 1.625 |
| Control | 0.00019 | 99 | 0.012 | 0.50 | 1.909 |

We sought to determine whether GRouNdGAN can generate samples from different cell types. If cell type information of the reference dataset is available a priori, one can easily generate realistic samples of each cell type separately (similar to the PBMC-CTL analysis) and use either cell type-specific GRNs, a shared GRN, or a combination of both. However, we asked whether our model could generate realistic samples *without* this knowledge, a more challenging task and useful for when such information is unavailable. Also, not requiring cell type information reduces the amount of user involvement (to annotate cell types) and improves model’s usability. We repeated the analysis using the PBMC-All dataset, containing eleven cell types. Although GRouNdGAN did not receive cell type/cluster information, we did not observe mode collapse (Figure S2), and it was able to generate cells from distinct clusters (Figure 2B, test set miLISI = 1.90). GRouNdGAN outperformed all simulators that did not use cell cluster information (Table 2) and compared to those that utilized this information, it was still the top performing based on MMD and RF AUROC and the second best according to the Euclidean distance (6% higher than the best) and miLISI (0.5% lower than the best) (Table 2). Finally, GRouNdGAN outperformed all other simulators based on all metrics when we repeated the analysis using the BoneMarrow dataset, containing continuous cell states and much fewer cells (Table S4). These results show that GRouNdGAN is a stable simulator of scRNA-seq data and even without extra cell type information in heterogenous datasets, it automatically generates realistic samples reflecting different cell types (while imposing a GRN). (See File S1 and Table S5 for stability analysis and the effect of different GRN properties on the performance).

**Table 2:** Performance of different simulators in generating realistic scRNA-seq data using the PBMC-All dataset. Baseline models that enable utilization of cell cluster labels or ratios are run with and without this information. The metrics are calculated between a simulated dataset of 1000 cells and the held-out test set of 1000 real cells. Each gene in the imposed GRN of GRouNdGAN is regulated by 15 TFs (constructed using GRNBoost2 from the experimental training set). For the first three metrics, a value closer to zero is preferred, for RF AUROC a value closer to 0.5 is preferred, and for miLISI a value closer to 2 is preferred. Best performance values (excluding control) are in bold-face. The control shows the value of the metrics comparing the two halves of the real dataset. For the first two metrics, the values correspond to the distance of the mean centroids of the real and simulated cells.

| Simulator | Cell cluster labels/ratios provided as input | Cosine distance | Euclidean distance | MMD | RF AUROC | miLISI |
| --- | --- | --- | --- | --- | --- | --- |
| GRouNdGAN | No | 0.00028 | 168 | **0.017** | **0.53** | 1.90 |
| cscGAN<sup>24</sup> | Yes | 0.00025 | 242 | 0.030 | 0.56 | **1.91** |
| cWGAN | Yes | 0.00053 | 239 | 0.027 | 0.57 | 1.89 |
| scGAN<sup>24</sup> | No | 0.00081 | 300 | 0.035 | 0.58 | 1.90 |
| scDESIGN2<sup>25</sup> | Yes | 0.00032 | 181 | 0.046 | 0.86 | 1.81 |
| scDESIGN2<sup>25</sup> | No | 0.00057 | 256 | 0.124 | 0.91 | 1.53 |
| SPARsim<sup>26</sup> | Yes | **0.00019** | **158** | 0.287 | 0.96 | 1.66 |
| SPARsim<sup>26</sup> | No | 0.00035 | 245 | 0.307 | 0.96 | 1.39 |
| Positive control | NA | 0.00022 | 116 | 0.012 | 0.50 | 1.91 |

### GRouNdGAN imposes a causal GRN in the simulated data

To assess the ability of GRouNdGAN in imposing the input causal GRN, we performed *in-silico* TF knockout experiments (one of GRouNdGAN’s capabilities) on the simulated cells from the PBMC-CTL dataset. We performed a forward pass after setting the expression of each TF to zero (one at a time) in the causal controller output, while keeping all other parameters and TF expressions unchanged, ensuring that the perturbations were performed on the same batch of cells forming matched case/control experiments. There was no change in the expression of genes that were not regulated by the knocked-out TF (as expected), while the expression of the regulated genes changed. Figure 2F shows the distribution of the adjusted p-values (Benjamini-Hochberg) for each TF-gene edge in the GRN, comparing the expression of a gene across all cells before and after the knockout of one of its regulating TFs (two-sided Wilcoxon signed-rank tests). In the majority (66.5%) of cases, the knockout of a gene’s regulating TF significantly (adjusted p-value < 0.05) altered its expression, showcasing that the GRN is indeed imposed in the expression profiles of simulated cells. Note that since each gene is regulated by multiple TFs, we cannot expect that knockout of a single TF result in a significant change in the expression of its target genes in all cases.

### The imposed GRN can be reconstructed from the simulated data

To test whether the imposed GRN can be reconstructed from the generated data, we first applied GRNBoost2^8^ to the reference PBMC-CTL dataset and recorded the top ten TFs for each gene. For each gene, we created two sets of five TFs based on their even/odd parity in the ranked list: TFs ranked 1^st^, 3^rd^, 5^th^, 7^th^, and 9^th^ were connected to the gene and were imposed (“positive control GRN”), while the remaining five TFs were not (“negative control GRN”), resulting in two GRNs with identical densities and the same number of TFs and target genes. When considering the positive and negative control GRNs as ground truths (separately), the GRN reconstructed using GRNBoost2 from the reference dataset resulted in comparable AUPRC (0.28 and 0.24, respectively, Table S6), reflecting that these edges are of comparable importance.

We simulated data using GRouNdGAN by imposing only the positive control GRN, and used GRNBoost2, PIDC^10^ GENIE3^9^, and PPCOR^37^ to reconstruct the underlying GRN (Figure 3, Figure S4, Table S6). All methods reconstructed the imposed edges with a much higher AUPRC compared to the reference dataset, while they did not assign high scores to the unimposed edges, resulting in close to random AUPRC. Figure 3 shows the performance of GRNBoost2 applied to the reference, GRouNdGAN-simulated, and scGAN-simulated data. The performance for scGAN (both GRNs) was lower than the reference dataset, showcasing that it had not preserved the TF-gene relationships of the experimental dataset well. The performance was relatively consistent between positive (Figures 3A and 3C) and negative control (Figures 3B and 3D) GRNs using the reference and scGAN-simulated data; however, it was much higher for the positive control GRN (compared to negative control GRN) using GRouNdGAN-simulated data. Similar patterns were observed when we repeated these analyses using the BoneMarrow dataset (Table S6, Figure S5). These results show that the imposed edges were accentuated by GRouNdGAN, while the unimposed edges were disrupted and could not be found by GRN inference methods.

**Figure 3:**
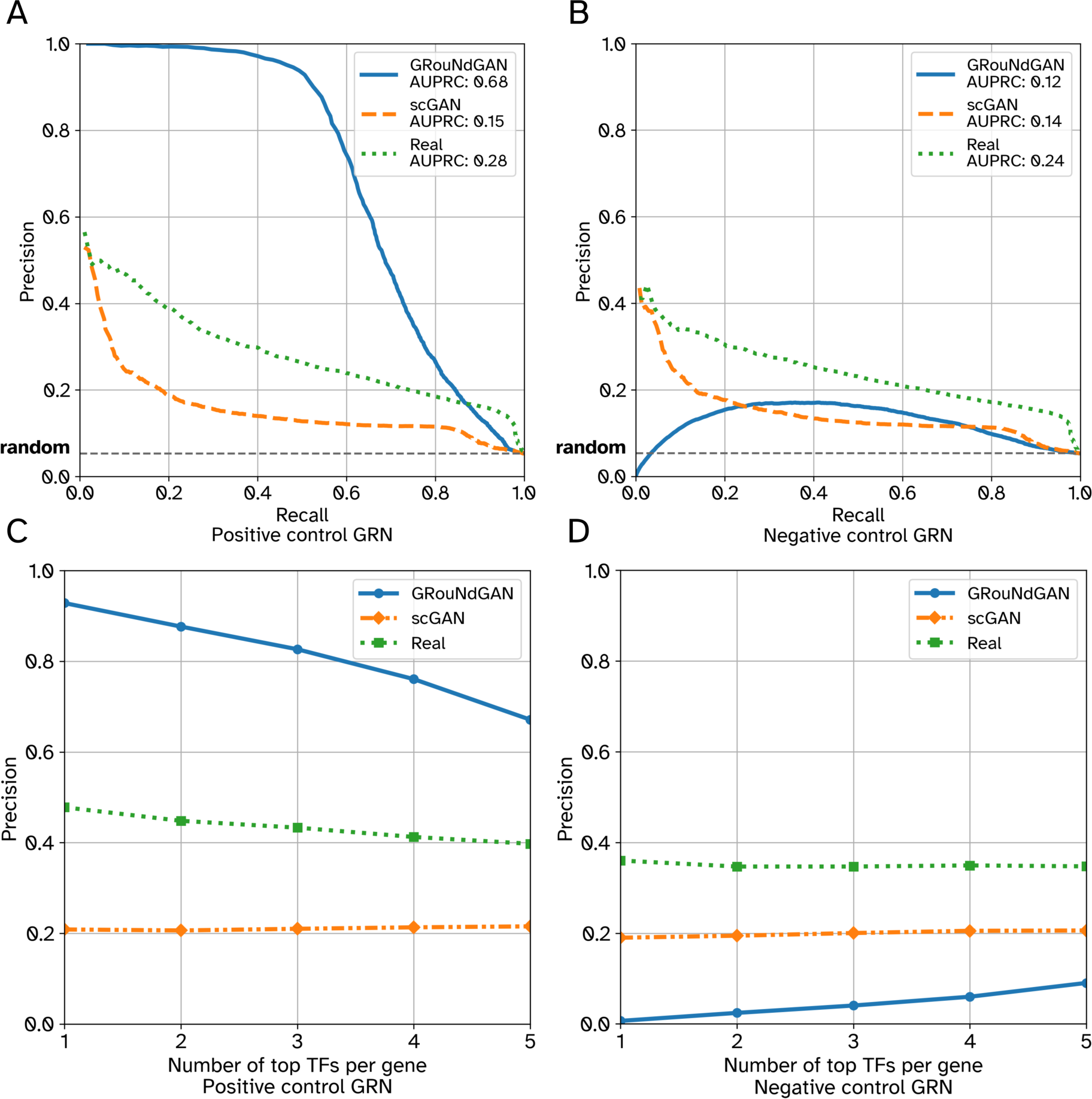
Performance of GRNBoost2 in recovering the imposed edges versus unimposed edges (PBMC-CTL). Left column shows the AUPRC (A) and Precision at k (per gene) (C) when the imposed edges (positive control GRN) were considered the ground truth. Right column shows the AUPRC (B) and Precision at k (per gene) (D) when the unimposed edges (negative control GRN) were considered the ground truth. Precision at k (per gene) refers to the precision when top k TFs for each gene is used to form the reconstructed GRN.

Next, we asked whether GRNBoost2 can reconstruct the imposed GRN, if GRNs of different densities are used to simulate data. Using data for which each gene was regulated by 15, 10, 5, and 3 TFs, we observed that in all cases the GRN imposed by GRouNdGAN could be inferred from the simulated data (Table S6). Finally, we repeated the controlled analysis above, swapping the role of positive and negative control GRNs. Similar to the results of Figure 3, the imposed edges were inferred by GRNBoost2, while the unimposed edges were not accurately discovered (Table S6). These results show that 1) GRouNdGAN imposes the causal GRN, 2) the GRN inference methods can identify the imposed edges, and 3) the TF-gene relationships present in the reference dataset but unimposed by GRouNdGAN are disrupted during simulation. The last property is particularly desirable for benchmarking GRN inference methods to ensure that the regulatory relationships present in the reference dataset (but unimposed) do not bias the simulated data, which could result in an inflated false positive rate.

### GRouNdGAN-simulated data preserves trajectories and can be used for pseudo-time inference

In addition to deciphering discrete states, scRNA-seq data is often used to determine continuous cell transitions during biological processes such as differentiation, using trajectory and pseudo-time inference^38–44^. We sought to determine whether GRouNdGAN-simulated data conforms to the transitional states and pseudo-time of the reference data. This is particularly important for benchmarking GRN inference methods that utilize pseudo-time information^12–14^. For this purpose, we used the BoneMarrow dataset corresponding to differentiation of hematopoietic stem cells to different lineages.

We used partition-based graph abstraction (PAGA)^38^ for trajectory inference of the reference and GRouNdGAN-simulated data. PAGA computes graph-like maps of data manifolds faithful to the topology of data, retaining its continuous and discrete structures. Figure S6 shows the PAGA graphs, where nodes capture discrete states, and the edges signify transitions between them. To compare the identified trajectories, we used markers of the cell types present in this dataset (Table S7)^38^. Figure 4 shows the expression patterns of two marker genes for erythroid cells (*Gata1, Klf1*), neutrophils (*Mpo, Ctsg*), and basophils (*Mcpt8, Prss34*) for PAGA trajectory graphs of both datasets (Figures S7-S13 shows marker genes of all cell types). The comparison showed similar activation patterns between corresponding regions of PAGA graphs of the experimental and GRouNdGAN data, revealing several trajectories (e.g., erythroid and neutrophil branches). Strikingly, even the number of nodes activated in each figure scaled similarly between the two datasets: e.g., the markers of basophils were consistently expressed in only one node in each dataset (Figures 4E-4F).

**Figure 4:**
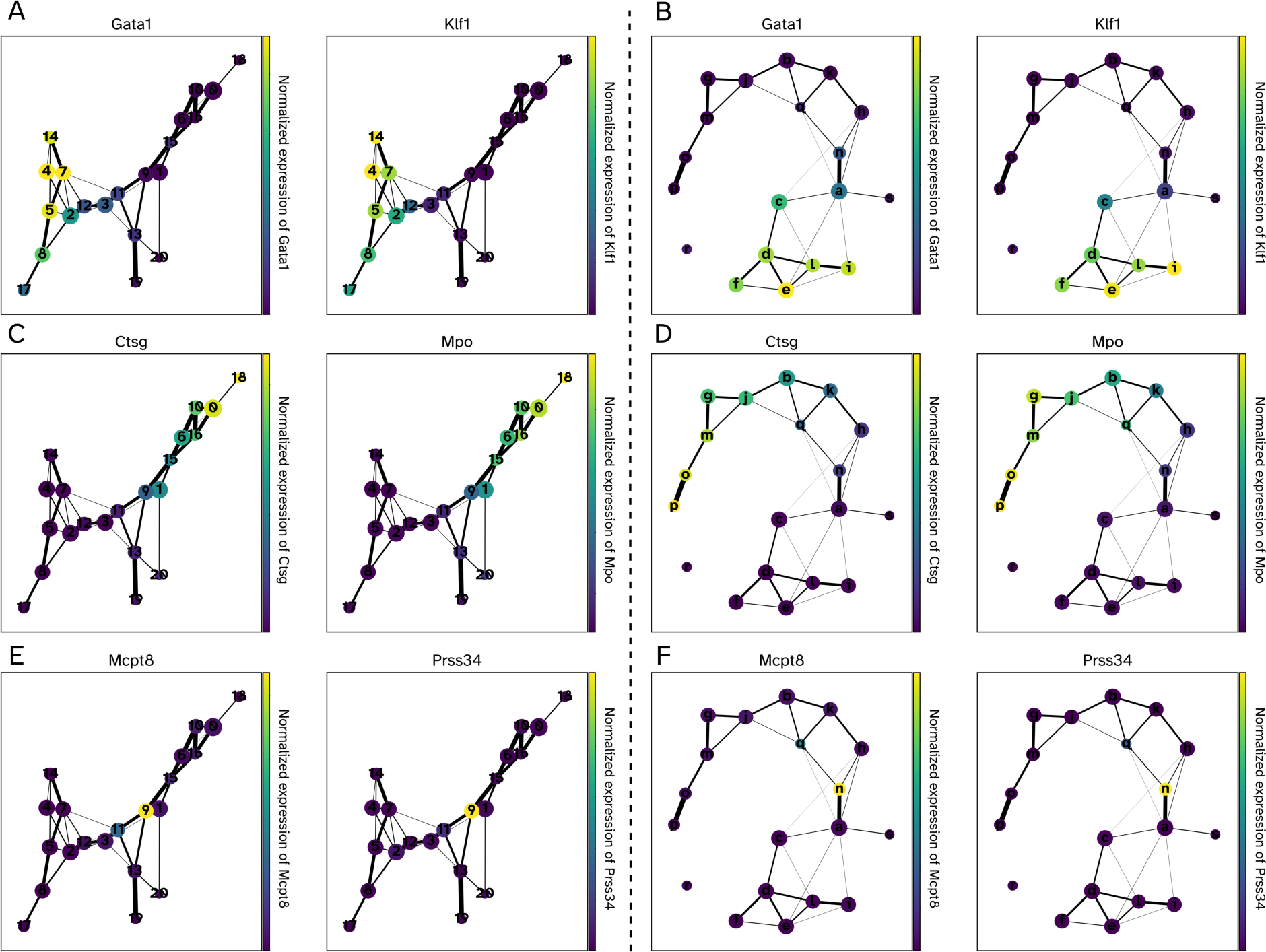
PAGA graphs and cell type marker gene activation patterns in experimental and simulated BoneMarrow dataset. In each figure, nodes represent Louvain clusters (capturing discrete states), edges represent transitions between states, and edge weights indicate confidence in the presence of connections. Low-connectivity edges below a threshold of 0.01 were discarded. Left panels (A, C, E) show the PAGA graph computed on GRouNdGAN-generated cells, and right panels (B, D, F) show the PAGA graph for the original BoneMarrow dataset. Node colors represent normalized expression of marker genes for erythroid cells (A, B), neutrophils (C, D), and basophils (E, F).

We annotated different cell types present in both datasets using these marker genes (following PAGA’s official tutorial) (Figures 5A-5B). The GRouNdGAN’s annotated graph showed similar topological features to that of the reference data and captured known biological properties of hematopoiesis, which were also noted by Wolf, et al.^38^ using the reference dataset (e.g., proximity between monocytes and neutrophils or association between megakaryocyte and erythroid progenitors). These observations show that the key characteristics of the reference data are retained in GRouNdGAN-simulated data.

**Figure 5:**
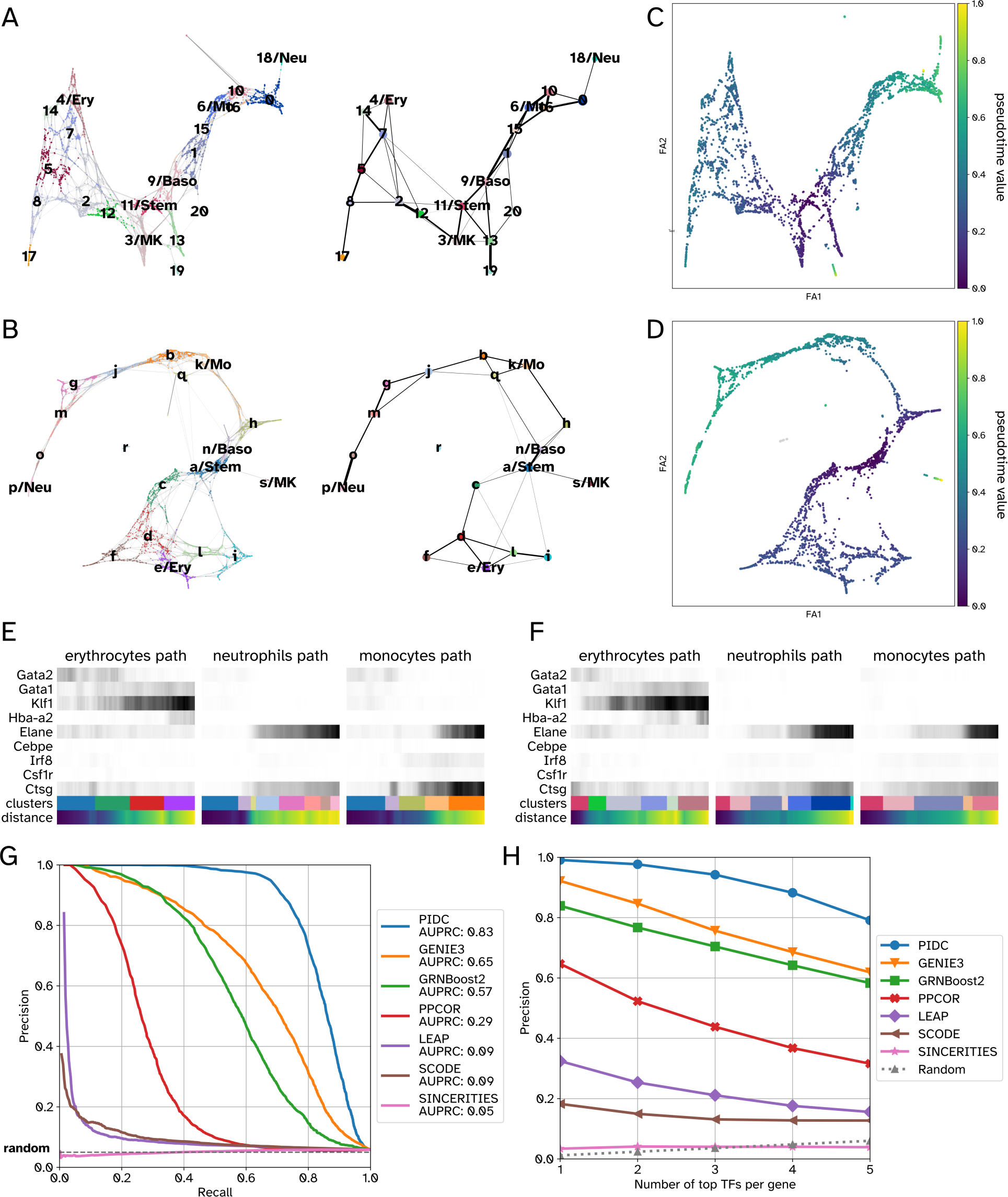
Consistency in trajectories and pseudo-temporal orderings between data generated by GRouNdGAN and the experimental BoneMarrow hematopoietic dataset. Panels A and B show the cell type annotations of simulated and experimental data, respectively. The figures on the left show the force-directed graphs (obtained using ForceAtlas2^45^) and the figures on the right show the PAGA graphs. Panels C and D show the PAGA-initialized force-directed graphs with cells colored by their inferred pseudo-time value. The following abbreviations are used: Stem for stem cells, Ery for erythroid cells, Neu for neutrophils, Mo for monocytes, MK for Megakaryocytes, and Baso for Basophils. Panels E and F display expression changes in the marker genes of erythrocyte (*Gata2*, *Gata1*, *Klf1*, and *Hba-a2*), neutrophil (*Elane* and *Cebpe*), and monocyte (*Irf8*, *Csf1r*, and *Ctsg*) branches for experimental (E) and simulated data (F). The distance refers to the geodesic distance from a root cell (here a stem cell). A darker shade of grey shows higher expression. Panels (G, H) show the performance of different GRN inference methods on data generated by GRouNdGAN using the BoneMarrow dataset.

We used the diffusion pseudotime algorithm^39^ (in scanpy^46^) to infer the progression of cells through geodesic distance (Figures 5C-5D). We noticed strong concordance when examining pseudo-time values assigned to cell subgroups in the reference and GRouNdGAN-generated data. To track these patterns, we focused on marker expression changes along erythroid, neutrophil, and monocyte trajectories (Figures S7-S9), similar to the analysis performed by Wolf, et al.^38^. In both datasets, the activation of neutrophils’ marker, *Elane,* and monocytes’ marker, *Irf8,* were predominantly observed towards the later stages of their trajectories (Figures 5E-5F). Additionally, we observed the activation of erythroid maturity marker genes *Gata2*, *Gata1*, *Klf1*, and *Hba-a2* roughly in sequential order along the erythroid trajectory in both datasets (Figures 5E-5F), concordant with previous findings^38^. The striking similarity between the activation patterns of these markers in both datasets and the preservation of their ordering highlights GRouNdGAN’s ability to capture dynamic transcriptional properties of scRNA-seq data, leading to correct trajectory inference and pseudo-time ordering.

### Benchmarking GRN inference methods using GRouNdGAN confirms prior insights from curated experimental datasets

The BEELINE study^19^ found conflicting results when benchmarking GRN inference methods on curated biological datasets and synthetic ones: top two performing models on synthetic data where among the bottom three on biological data. We re-investigated this using GRouNdGAN-simulated data with the BoneMarrow dataset and the positive and negative control GRNs (described earlier). We used seven GRN inference methods used by BEELINE, GRNBoost2^8^, GENIE3^9^, PIDC^10^, PPCOR^37^, LEAP^12^, SCODE^13^, and SINCERITIES^14^, capturing a wide range of methods with a broad spread of performances reported on synthetic and curated data in the BEELINE study. We observed a striking resemblance between the performance pattern (order) of the methods on GRouNdGAN-simulated data and curated biological data from BEELINE (Figure 5G-5H and Table S6 versus Figure 4 of BEELINE^19^): the only difference was the swapping of LEAP and SCODE’s order. We repeated this analysis using the PBMC-CTL dataset (Table S6), obtaining similar patterns.

Overall, PIDC^10^ outperformed all methods using both datasets, followed by GENIE3^9^ and GRNBoost2^8^. LEAP^12^, SCODE^13^ and SINCERITIES^14^ (which use pseudo-time and could only be applied to the BoneMarrow dataset) performed worse than others, matching their behavior in the BEELINE study on curated biological data (but not on synthetic data). All methods performed close to random on unimposed negative control GRN edges. These results not only show that GRouNdGAN can be used for benchmarking GRN inference methods, but also shows that the insights obtained from it matches those obtained from curated biological datasets, the formation of which requires extensive amount of resources and effort.

### *In-silico* perturbation experiments using GRouNdGAN

In many studies, one is interested to characterize the relationship between TFs’ expression and phenotypic labels (e.g., cell types). Differential expression (DE) analysis is a common approach to identify TFs (or other genes) most associated with a specific cell type. However, to directly test whether perturbing the expression of such candidate TFs change the expression of genes representative of a cell type, perturbation experiments (e.g., TF knockout) are desired. As a causal implicit generative model, GRouNdGAN provides the capability to sample from interventional distributions, thereby enabling *in-silico* perturbation experiments. This functionality is provided at inference time by enabling manipulation of TFs’ expressions as the outputs of the causal controller. This change is propagated along the GRN (through the model architecture), impacting the expression of genes regulated by the perturbed TF(s). By enabling a deterministic mode of operation, GRouNdGAN allows for exact comparison of gene expressions before and after perturbation while maintaining the invariance of other parameters (Methods).

To test this functionality, we asked whether the knockout of top 3 TFs most differentially expressed (Mann Whitney U test) between each cell type and the rest (identified from the reference dataset) would result in less cells being generated in the vicinity of the cells of that type. If positive, this implies that the perturbation in the TFs’ expression has modified the gene expression profile of generated cells so that they no longer resemble that particular cell type (confirming our expectation). Figure 6A shows the UMAP embedding of the reference dataset (miLISI = 1.94 when compared with simulated data) and Figure 6B shows the density plots of the experimental and simulated data, revealing a high degree of resemblance. We focused on the changes in the iLISI values of experimentally profiled cells of a specific type (annotated in the original study), when jointly mapped to the same embedding space with simulated data, before and after knockout. For the changes in the iLISI values to be observable, it was necessary to focus on cell types that occupy relatively distinct regions of the embedding space: CD14+ monocytes, CD19+ B cells, Dendritic cells, CD56+ natural killer cells (Figure S14); otherwise, the presence of other cell types in that region would confound the results. Figure 6C shows the iLISI distributions for each cell type (only for real cells), calculated along with unperturbed simulated cells (blue) and perturbed simulated cells (orange). Figure 6D-6E show the iLISI values and the distribution of generated cells after the knockout experiment for CD19+ B cells as an example (see Figures S15-S18 for other cell types). In all cases, the iLISI values significantly reduced after the knockout experiment (Figure 6C, Table S8), showcasing that after the knockout of top TFs associated with a cell type, GrouNdGAN generated much fewer cells resembling that cell type (even though it did not have the knowledge of the cell type annotations). Instead, the simulated cells were dispersed into other cell types, yet they retained meaningful positions within the overall dataset embedding space. Also, the miLISI of the other cell types remained relatively unchanged (Table S8). This analysis shows that *in-silico* TF perturbation experiments using GrouNdGAN produces results concordant with results directly obtained from real biological data, a direction that we will explore further in the future studies.

**Figure 6:**
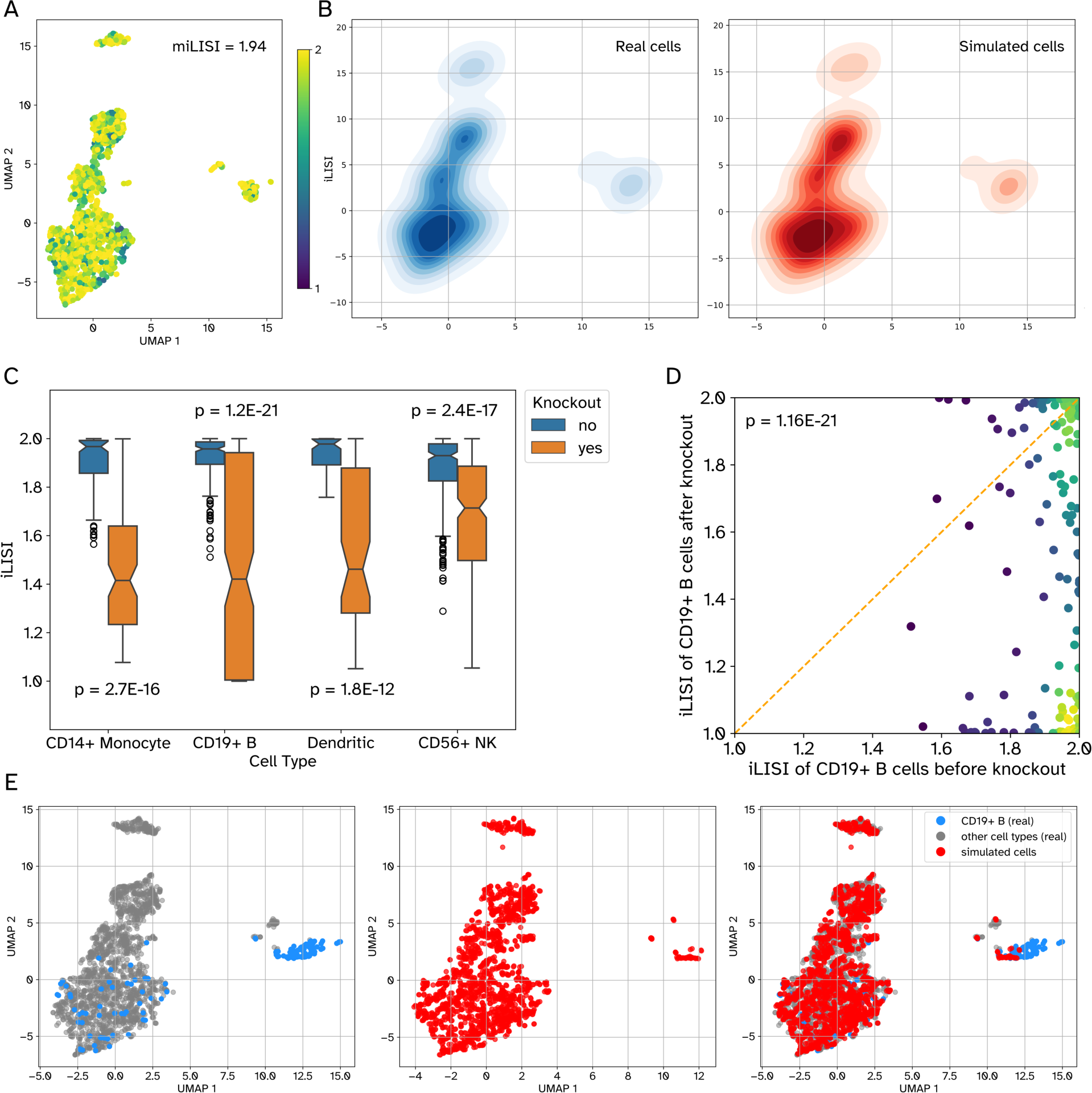
TF knockout experiments and their effect on cell type generation. A) The UMAP shows the distribution of 2000 randomly selected cells from the experimental PBMC-All dataset. The color shows the iLISI value of each cell (miLISI = 1.94) calculated from a UMAP embedding, jointly obtained from the experimental cells and the same number of GRouNdGAN-generated cells. TFs were omitted as features when generating UMAP plots. In the simulated data, each gene was regulated by 15 TFs, identified using GRNBoost2. B) Each plot shows the density of cells in the embedding space: the left plot (blue) corresponds to the real cells and the right plot (red) shows the simulated cells. C) The boxplots show the distribution of iLISI values of real cells, calculated along with unperturbed simulated cells (blue) and with perturbed simulated cells (orange). For each cell type, top three most differentially expressed TFs were knocked out. The p-values reported in this figure were calculated using one-sided Wilcoxon signed rank tests. D) The scatter plot shows the iLISI values of CD19+ B cells calculated along with unperturbed simulated cells (x-axis) and along with perturbed simulated cells (y-axis). The circles correspond to real cells and their colors reflect the density of datapoints in that region. E) The UMAP embedding of real cells (left), simulated cells after knockout of top TFs of CD19+ B cells (middle), and all together (right).

## Discussion

GRouNdGAN is a causal implicit generative model designed to simulate realistic scRNA-seq datasets based on a user-defined GRN and a reference dataset. By incorporating GRN connections in its architecture and including auxiliary tasks, it imposes causal TF-gene relationships in the gene expression profile of simulated cells. These causal relationships are verifiable by *in-silico* knockout experiments and identifiable by GRN inference methods. (In addition to the experimental analyses performed in this study, an interested reader should refer to Kocaoglu et al.^31^ for theoretical evidence of causality of data generated by this architecture.)

We demonstrated that GRouNdGAN achieves state-of-the-art performance on various tasks including realistic scRNA-seq data generation, GRN inference benchmarking, and *in-silico* knockout experiments. Even when applied to heterogenous datasets, it generated realistic datasets without utilizing information regarding cell clusters, achieving best or second-best performance compared to models that used this information. We should note that it is trivially possible to use cell type/cluster information with GRouNdGAN: one can simply provide subsets of the experimental dataset corresponding to a cell type/cluster as a reference, one at a time with a cell-type specific (or shared) GRN. However, the fact that this model does not require this information broadens its applicability. We showed that GRouNdGAN-generated data preserves complex cellular dynamics and patterns of the reference dataset such as lineage trajectories, pseudo-time orderings, and patterns of cell type markers’ activation.

GRouNdGAN imposes user-defined causal regulatory interactions, while excluding those present in the reference dataset, but not in the input GRN. This makes GRouNdGAN-simulated datasets ideal for benchmarking GRN inference algorithms with the input GRN as the ground truth. We benchmarked seven popular GRN inference algorithms and showed that our results coincide with results reported by BEELINE^19^ on experimental datasets. This highlights GRouNdGAN’s ability to bridge the gap between experimental and simulated datasets for GRN benchmarking without requiring excessive user manipulation. Additionally, since GRouNdGAN generates data based on a structural causal model, it can be used to sample synthetic cells from both observational and interventional distributions. Using this property, we conducted *in-silico* perturbation experiments by knocking out key regulating TFs in various cell types and observed that the targeted cell-states were suppressed in the generated dataset.

GRouNdGAN maintains key advantages over existing simulators that incorporate GRNs such as SERGIO and BoolODE. During training, GRouNdGAN’s parameters are optimized to minimize any mismatch between real and simulated data distributions. This enables it to replicate scRNA-seq data characteristics (e.g., technical and biological noise) without any user intervention or post-processing. In contrast, SERGIO simulates data by first generating a “clean” dataset based on SDEs. Then, the user would iteratively fine-tune three noise parameters until the simulated and reference datasets are matched. This data-matching procedure puts extra burden on the user and might lead the user to overly rely on fine-tuning technical noise parameters, instead of SDE parameters. Overreliance on added technical noise to achieve data-matching can potentially corrupt the data, rendering the underlying regulatory patterns unobservable in the final simulated dataset. This might explain the near-random GRN inference performance reported by SERGIO on their noisy datasets^4^. GRouNdGAN is trained such that each of its generated genes or TFs mimics the expression patterns of a corresponding gene or TF in the reference dataset, preserving gene/TF identities. In contrast, BoolODE and SERGIO simulate pseudo-genes whose identities cannot be readily mapped to their counterparts in the reference dataset. Gene identity preservation can be attained in these simulators by fine-tuning the SDE parameters until the expression patterns of a given generated gene aligns with a corresponding gene in the reference dataset. This, however, places further burden on the user. Finally, GRouNdGAN learns complex non-linear co-regulatory patterns from the data, instead of relying on simplifying assumptions or user input. In contrast, SERGIO assumes that the combined effect of multiple regulatory TFs is simply the sum of their individual effects. Similarly, BoolODE requires the user to provide a truth table specifying combinatory TF co-regulation rules for each target gene. In contrast, GRouNdGAN’s neural network-based generator enables it to implicitly learn complex non-linear co-regulatory patterns that may represent co-expression, co-inhibition, or co-activation of individual genes. This is achieved during the training of GRouNdGAN without user intervention.

Several key points need to be considered when using GRouNdGAN. First, our analyses showed that the architectural and hyperparameter choices of the presented model result in good performance across different datasets. We performed hyperparameter tuning on one dataset and used the same set of values for all three datasets and for different choices of GRNs, obtaining state-of-the-art performance. However, when trained on a completely new dataset, it is still advisable to test different choices of the hyperparameters using a validation set. Second, the default architecture of the model includes LSN layers and, as such, the model generates library-size normalized expression values. Our ablation study (File S1) showed that without the LSN layer, GRouNdGAN can still generate realistic profiles; however, including it improves the GRN imposition and stabilizes the training. Third, an important consideration for any causal inference problem is that given an observational dataset, its underlying causal graph is not unique. When benchmarking GRN inference methods, this problem exists regardless of how the ground truth GRN is obtained (even using knockout/knockdown experiments), and a GRN inference algorithm may find a plausible causal GRN, yet not the one that generated the data. Despite this, our analysis using the positive/negative control GRNs showed that GRN inference methods could detect the imposed edges while ignoring others that were unimposed. Based on this observation, we posit that the imposed GRN by GRouNdGAN, while not the only causal GRN capable of generating the simulated dataset, is the most probable one. However, verifying this requires theoretical analyses that are well beyond the scope of this study.

Finally, although GRouNdGAN can generate data with any GRN, the quality of the data deteriorates if the imposed TF-gene relationships are significantly different from those consistent with the patterns in the reference dataset. The reason is that in such a scenario, generating realistic simulated datapoints and imposing the GRN act as contradictory requirements. This is a fundamental problem that is not unique to GRouNdGAN and is applicable to any simulator with these two requirements. However, simulators that do not try to impose a causal graph (e.g., scGAN) can focus on generating realistic scRNA-seq samples, even at the expense of disrupting some of the interactions present in the reference dataset (as we observed in Figure 3 with scGAN). As a result, for GRN inference method benchmarking, we recommend first finding a set of regulatory edges from the reference dataset (e.g., using GRNBoost2) and imposing a selected subset of those in the simulated data. This ensures that the imposed edges are encoded in the simulated data, the unimposed (or undetected) edges are disrupted (as shown in our analyses), and the simulated dataset resembles the reference dataset. It is important to note that the regulatory edges inferred from the reference dataset do not need to be all causal; some of them could represent spurious correlations that are observable in the data. In fact, most GRN inference methods do not identify causal edges, but can be used to form the input GRN. When a subset of the (potentially) non-causal edges of the reference dataset are imposed by GRouNdGAN, they are imposed in a causal manner, representing the *causal* graph underlying the *simulated* data (and not the reference data).

The current version of GRouNdGAN only allows imposing a bipartite GRN and does not support regulation of TFs by other TFs, an assumption that we plan to relax with the future versions of the model. Moreover, we intend to augment the model’s architecture in the future to enable imposing a causal graph with three layers of nodes capturing TF-gene-phenotype relationships. This is particularly useful for development and evaluation of phenotype-relevant GRNs, a concept that we have introduced in the past^11^. Additionally, a conditional version of GRouNdGAN can be developed, allowing the user to generate cells of specific cell type or on a specific stage along a lineage trajectory. Finally, by incorporating DAG structural learning^47^, we intend to enable GRouNdGAN to infer a GRN while simultaneously learning to mimic the reference dataset.

## Methods

### Datasets and preprocessing

We downloaded the human peripheral blood mononuclear cell (PBMC Donor A) dataset containing the single-cell gene expression profiles of 68579 PBMCs represented by UMI counts from the 10x Genomics (https://support.10xgenomics.com/single-cell-gene-expression/datasets/1.1.0/fresh_68k_pbmc_donor_a). This dataset contains a large number of well-annotated cells and has been used by other models to generate synthetic scRNA-seq samples^24^. We also downloaded the scRNA-seq profile of 2730 cells corresponding to differentiation of hematopoietic stem cells to different lineages from mouse bone marrow^34^ (“BoneMarrow” dataset) from Gene Expression Omnibus (accession number: GSE72857)

We followed the pre-processing steps of scGAN and cscGAN^24^ using scanpy version 1.8.2^46^. In each dataset, cells with nonzero counts in less than ten genes were removed. Similarly, genes that only had nonzero counts in less than three cells were discarded. Next, library-size normalization was performed on the counts per cell with a library size equal to 20,000 in order to be consistent with previous studies ^24^. Next, top 1000 highly variable genes were selected using the dispersion-based method described by Satija et al.^48^.

### GRouNdGAN’s model architecture

GRouNdGAN consists of 5 components, each implemented using separately parameterized neural networks: a causal controller, target generators, a critic, a labeler, and an anti-labeler (Figure 1B).

#### Causal controller

The role of causal controller is to generate the expression of TFs that causally control the expression of their target genes based on a user-defined gene regulatory network (Figure 1B). To achieve this, it is first pre-trained (Figure 1A) as the generator of a Wasserstein GAN (WGAN). GANs are a class of deep learning models that can learn to simulate non-parametric distributions^49^. They typically involve simultaneously training a generative model (called the “generator”) that produces new samples from noise, and its adversary, a discriminative model that tries to distinguish between real and generated samples (called the “discriminator”). The generator’s goal is to generate samples so realistic that the discriminator cannot determine whether it is real or simulated (an accuracy value close to 0.5). Through adversarial training, the generator and discriminator receive feedback, allowing them to co-evolve in a symbiotic manner.

The main difference between a WGAN and a traditional GAN is that in the former, a Wasserstein distance is used to quantify the similarity between the probability distribution of the real data and the generator’s produced data (instead of Kullback–Leibler or Jensen–Shannon divergences). Wasserstein distance has been shown to stabilize the training of WGAN without causing mode collapse^32^. The detailed formulation of the Wasserstein distance used as the loss function in this study is provided in File S1. In addition, instead of a discriminator, WGAN uses a “critic” that estimates the Wasserstein distance between real and generated data distributions. In our model, we added a gradient penalty term for the critic (proposed by Gulrajani et al.^50^ as an alternative to weight clipping used in the original WGAN) in order to overcome vanishing/exploding gradients and capacity underuse issues.

In the pretraining step, we trained a WGAN with a generator (containing an input layer, three hidden layers, and an output layer), a library-size normalization (LSN) layer^24^, and a critic (containing an input layer, three hidden layers, and an output node). A noise vector of length 128, with independent and identically distributed elements following a standard Gaussian distribution, was used as the input to the generator. The output of the generator was then fed into the LSN layer to generate the gene and TF expression values. The details of hyperparameters and architectural choices of this WGAN are provided in Table S1 (in File S1). Although we were only interested in generating expression of TFs using the generator of this WGAN (in the second step of pipeline), the model was trained using all genes and TFs to properly enforce the library-size normalization. Once trained, we discarded the critic and the LSN layer, froze the weights of the generator and used it as the “causal controller”^31^ to generate expression of TFs (Figure 1B).

#### Target generators

The role of target generators is to generate the expression of genes causally regulated by TFs based on the topology of a GRN. Consider a target gene *Gj* regulated by a set of *TFs*: {*TF*_1_, *TF*_2_, …, *TF_n_*}. Under the causal sufficiency assumption and as a result of the manner by which TFs’ expressions are generated from independent noise variables, we can write *E_Gj_* = *f_Gj_*(*E_TF_*_1_, *E_TF_*_2_, …, *E_TFn_*, *N_Gj_*) and *E_TFi_* = *f_TFi_*(*N_TFi_*) for *i* = 1, 2, …, *n*, where *E* represents expression, *N* represents a noise variable, and *f* represents a function (to be approximated using neural networks). All noise variables are jointly independent. Following the theoretical and empirical results of CausalGAN^31^, we can use feedforward neural networks to represent functions *f* by making the generator inherit its neural network connections from the causal GRN. To achieve this, we generate each gene in the GRN by a separate generator such that target gene generators do not share any neural connections (Figure 1B).

As input, the generator of each gene accepts a vector formed by concatenating a noise variable and a vector of non-library size normalized TF expressions from the causal controller, corresponding to the TFs that regulate the gene in the imposed GRN (i.e., its parents in the graph). The expression values of TFs and the generated values of target genes are arranged into a vector, which is passed to an LSN layer for normalization. We used target generators with three hidden layers of equal width. The width of the hidden layers of a generator is dynamically set as twice the number of its regulators (the noise variable and the set of regulating TFs). If the imposed GRN is relatively dense and contains more than 5000 edges, we set the depth to 2 and the width multiplier to 1 to be able to train on a single GPU. The details of hyperparameters and architectural choices are provided in Table S2 (in File S1).

In practice, generating each target gene’s expression using separate neural networks introduces excess overhead. This is because in every forward pass, all target genes’ expressions must be first generated before collectively being sent into the LSN layer. As a result, instead of parallelizing the generation of each target gene’s expression (which due to the bottleneck above does not provide a significant computational benefit), we implemented target generators using a single large sparse network. This allows us to reduce the overhead and the training time to train the model on a single GPU, and to benefit from GPU’s large matrix multiplication. We mask weights and gradients to follow the causal graph, while keeping the generation of genes independent from each other. From a logical standpoint, our implementation has the same architecture described earlier, but is significantly more computationally efficient.

#### Critic

Similar to a traditional WGAN, the objective of GRouNdGAN’s critic (Figure 1B) is to estimate the Wasserstein distance between real and generated data distributions. We used the same critic architecture as the WGAN trained in the first stage.

#### Labeler and anti-labeler

Although the main role of the target generators is to produce realistic cells to confuse the critic, it is crucial that they rely on the TFs’ expression (in addition to noise) in doing so. One potential risk is that the target generators disregard the expression of TFs and solely rely on the noise variables. This is particularly probable when the imposed GRN does not conform to the underlying gene expression programs of real cells in the reference dataset; in such a scenario, and to make realistically looking simulated cells, it is more convenient for a WGAN to simply ignore the strict constraints of the GRN and solely rely on noise.

To overcome this issue, we used auxiliary tasks and neural networks known as “labeler” and “anti-labeler”^31^. The task of these two networks is to estimate the causal controller’s TF expressions (here called labels) from the target genes’ expressions alone (by minimizing the squared L2 norm between each element’s TF estimates and their true value). This resembles the idea behind an autoencoder and ensures that the model will not disregard the expression of TFs in generating the expression of their target genes. The anti-labeler is trained solely based on the outputs of the target generators, while the labeler utilizes both the outputs of target generators and the expression of real cells (Figure 1B). They are both implemented as fully connected networks with a width of 2000 and a depth of 3 and optimized using the AMSGrad^51^ algorithm. Each layer, except the last one, utilizes a ReLU activation function and batch normalization. In addition to WGAN losses, we add labeler and anti-labeler losses to the generator to minimize both. This is different from the approach used in CausalGAN, where the anti-labeler’s loss is maximized in the early training stages to ensure that the generator doesn’t fall into label-conditioned mode collapse. In GRouNdGAN, the causal controller is pretrained and generates continuous labels (TFs expression) and does not face a similar issue. As a result, we instead minimized the loss of the anti-labeler from the beginning and as such the labeler and anti-labeler both act as auxiliary tasks to ensure the generated gene expression values take advantage of TFs expression.

### Training procedure and hyperparameter tuning

We follow a two-step training procedure comprising of training two separate WGANs (Figure 1) to train GRouNdGAN. The generators and critics in both GANs are implemented as fully connected neural networks with rectified linear unit (ReLU) activation functions in each layer, except for the last layer of the critic. The weights were initialized using He initialization^52^ for layers containing ReLU activation and Xavier initialization^53^ for other layers (containing linear activations). We used Batch Normalization^54^ to normalize layer inputs for each training minibatch, except for the critic. This is since using it in the critic invalidates the gradient penalty’s objective, as it penalizes the norm of the critic’s gradient with respect to the entire batch rather than to inputs independently. An LSN layer^24^ was used in both WGANs (Figure 1A and 1B) to scale counts in each simulated cell to make it consistent with the library size of input reference dataset. This normalization results in a dramatic decrease in convergence time and smooths training by mitigating the inherent heterogeneity of scRNA-seq data.

GRouNdGAN solves a min-max game between the generator (*f_g_*) and the critic (*f_c_*), with the following objective function:

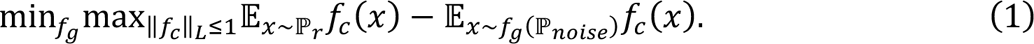

We alternated between minimizing the generator loss for one iteration and maximizing the generator loss for five iterations. We employed the AMSGrad^51^ optimizer with the weight decay parameters *β*_1_ = 0.5, *β*_2_ = 0.9 and employed an exponentially decaying learning rate for the optimizer of both the critic and generator.

The hyperparameters were tuned using a validation set consisting of 1000 cells from the PBMC-CTL dataset (Tables S1-S2 in File S1) based on the Euclidean distance and the RF AUROC score, which were consistently in accord. The same hyperparameters were used for all other analyses and the other two datasets (PBMC-All and BoneMarrow).

### Causal GRN preparation

This section describes the creation of the causal graph inputted to GRouNdGAN used to impose a causal structure on the model. GRouNdGAN accepts a GRN in the form of a bipartite directed acyclic graph (DAG) as input, representing the relationship between TFs and their target genes. In this study, we created the causal graph using the 1000 most highly variable genes identified in the preprocessing step. First, the set of TFs among the highly variable genes were identified based on the AnimalTFDB3.0 database^55^ and a GRN was inferred using GRNBoost2^8^ (with the list of TFs provided). Table S5 shows the characteristics of the different GRNs used in this study. It is important to note that the regulatory edges identified from the reference dataset using GRNBoost2 are not necessarily causal edges (and they do not need to be for the purpose of forming the input GRN), but they are consistent with the patterns of the data. However, when this (potentially non-causal) GRN is imposed by GRouNdGAN, it is imposed in a causal manner and represents the causal data generating graph of the simulated data (and not the reference data).

### Evaluation of the resemblance of real and simulated cells

We evaluated all models using held-out test sets containing randomly selected cells from each reference dataset (1000 cells from PBMC-CTL and PBMC-All and 500 cells from BoneMarrow). To quantify the similarity between real and generated cells, we employed various metrics. For each cell represented as a datapoint in a low dimensional embedding (e.g., t-SNE or UMAP), the integration local inverse Simpson’s Index (iLISI)^36^ captures the effective number of datatypes (real or simulated) to which datapoints of its local neighborhood belong based on weights from a Gaussian kernel-based distributions of neighborhoods. The miLISI is the mean of all these scores and in our study ranges between 1 (poor mixing of real and simulated cells) and 2 (perfect mixing of real and simulated cells). Additionally, we calculated the cosine and Euclidean distances of the centroids of real cells and simulated cells, where the centroid was obtained by calculating the mean along the gene axis (across all simulated or real cells).

To estimate the proximity of high-dimensional distributions of real and simulated cells without creating centroids, we used the maximum mean discrepancy (MMD)^35^. Given two probability distributions *p* and *q* and a set of independently and identically distributed (i.i.d.) samples from them, denoted by *X* and *Y*, MMD with respect to a function class ℱ is defined as

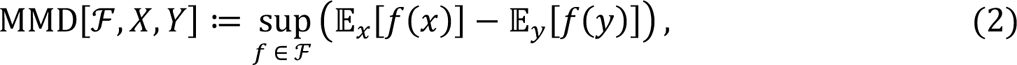

where sup refers to supremum and 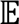 denotes expectation. When the MMD function class ℱ is a unit ball in a reproducing kernel Hilbert space (RKHS) ℋ with kernel *k*, the population MMD takes a zero value if and only if *p* = *q* and a positive unique value if *p* ≠ *q*. The squared MMD can be written as the distance of mean embeddings *μ_p_*, *μ_q_* of distributions *p* and *q*, which can be expressed in terms of kernel functions:

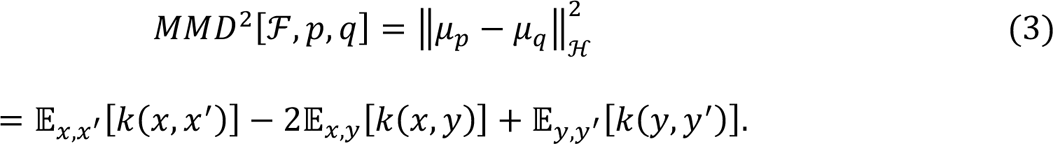

Following existing implementations of MMD in the single-cell domain^24, 56^, we chose a kernel that is the sum of three Gaussian kernels to increase sensitivity of the kernel to a wider range:

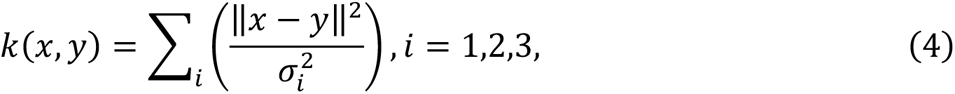

where σ*_i_* denote standard deviations and were chosen to be the median of the average distance between a point to its 25 nearest neighbors divided by factors of 0.5, 1, and 2 in the three kernels, respectively.

We also used a random forests (RF) classifier and used its area under the receiver operating characteristic (AUROC) curve to determine whether the real and simulated cells can be distinguished from each other. Consistent with previous studies^24, 57^, we first performed a dimensionality reduction using principal component analysis (PCA) and used the top 50 PCs of each cell as the input features to the RF model, which improves the computational efficiency of this analysis. The RF model was composed of 1000 trees and the Gini impurity was used to measure the quality of a split.

### Baseline simulator models

We compared the performance of GRouNdGAN to scDESIGN2^25^, SPARsim^26^, and three GAN-based methods: scGAN^24^, cscGAN with projection-based conditioning^24^, and a conditional WGAN (cWGAN). The cWGAN method conditions by concatenation following the cGAN framework^58^. More specifically, it concatenates a one-hot encoded vector (representing the cluster number or cell type) to the noise vector input to the generator and cells forwarded to the discriminator. We did not train the cWGAN or cscGAN on the PBMC-CTL dataset, since it contains only one cell type. For the PBMC-All and the BoneMarrow dataset, we trained all models above. Additionally, we simulated data using scDESIGN2 and SPARsim with and without cell cluster information, as they allow providing such side information in their training.

To train models that utilized cell cluster information, we performed Louvain clustering and provided the cluster information and ratio of cells per cluster during training. Clustering was done by following the cell ranger pipeline^33^, based on the raw unprocessed dataset (and independent of the pre-processing steps described earlier for training simulators). First, genes with no UMI count in any of the cells were removed. Then the gene expression profile of each cell was normalized by the total UMI of all (remaining) genes, and highly variable genes were identified. The gene expression profile of each cell was then re-normalized by the total UMI of retained highly variable genes, and each gene vector (representing its expression across different cells) was z-score normalized. Given the normalized gene expression matrix above, we found top 50 principal components (PCs) using PCA analysis. These PCs were then used to compute a neighborhood graph with a local neighborhood size of 15, which was used in Louvain clustering. We ran the Louvain algorithm with a resolution of 0.15.

For SPARSim, we set all sample library sizes to 20000 and estimated gene expression level intensities and simulation parameters by providing it with both raw and normalized count matrices. When cell cluster information was provided, distinct SPARSim simulation parameters were estimated per cell for each cluster. scDESIGN2 accepts input matrices where entries are integer count values; we thus performed rounding on the expression matrix before fitting scDESIGN2. With cluster information provided, a scDESIGN2 model was fit separately for cells of each cluster, and similar to conditional GANs, the ratio of cells per cluster was provided to the method.

### *In-silico* perturbation experiments using GroundGAN

To perform perturbation experiments using GroundGAN, we put the trained model in a deterministic mode of operation. This is necessary to ensure that the perturbation experiments are performed on the same batch (i.e., replicate) of generated cells to form matched case/control experiments. To do this, we performed a forward pass through the generator and then saved the input noise to the causal controller, the input noise to the target generators, the TF expression values generated by the causal controller, and the per-cell scaling factor of the LSN layer. Subsequent passes through the generators used the saved parameters so that ensuing runs always output the same batch of cells (instead of generating new unmatched cells).

### Trajectory inference and pseudo-time analysis

Following official PAGA tutorial for the BoneMarrow dataset (https://github.com/scverse/scanpy-tutorials/blob/master/paga-paul15.ipynb), we used (Partition-based graph abstraction) PAGA^38^ for trajectory inference and analysis. We built force-directed graphs^59^ (with ForceAtlas2^45^) using the top 20 principal components of the data (using principal component analysis or PCA) and a neighborhood graph of observations computed using UMAP (to estimate connectivities). We next denoised the graph by representing it in the diffusion map space and computed distances and neighbors as before using this new representation. After denoising, we then ran the Louvain clustering algorithm with a resolution of 0.6. Finally, we ran the PAGA algorithm on the identified clusters and used the obtained graph to initialize and rebuild the force-directed graph.

### GRN inference methods

In our GRN benchmarking analysis, we focused on seven GRN inference algorithms: GENIE3^9^, GRNBoost2^8^, PPCOR^37^, PIDC^10^, LEAP^12^, SCODE^13^ and SINCERITIES^14^, which were used in the BEELINE study^19^. Of these methods, LEAP requires pseudo-time ordering of cells, while SCODE and SINCERITIES require both pseudo-time ordering and pseudo-time values. Since not all algorithms inferred the edge directionality or its sign (activatory or inhibitory nature), we did not consider these factors in our analysis to be consistent among different models. We ran these models as docker containers using the docker images provided by BEELINE’s GitHub (https://github.com/Murali-group/Beeline)^19^ with the default parameters used in BEELINE. To benchmark algorithms requiring pseudo-time ordering of cells, we computed the pseudo-times of GRouNdGAN-simulated data (based on the BoneMarrow dataset) using PAGA^38^ and diffusion pseudotime^39^, following the methodology described earlier. In the GRN inference benchmark analysis, we did not provide the list of TFs to GRNBoost2 to make it consistent with other GRN inference methods.

## Authors’ contributions

All authors contributed to the design of the study and the algorithm, the analyses, and the writing of the manuscript. AE supervised the study. YZ implemented the model and performed the statistical analyses. All authors read and approved the final manuscript.

## Acknowledgements

This work was supported by grants from Natural Sciences and Engineering Research Council of Canada (NSERC) [RGPIN-2019-04460] (AE), Government of Canada’s New Frontiers in Research Fund (NFRF) [NFRFE-2019-01290] (AE), Canada Foundation for Innovation (CFI) JELF [project 40781], and McGill Initiative in Computational Medicine (AE). This research was enabled in part by support provided by the Digital Research Alliance of Canada (https://alliancecan.ca) and Compute Canada (www.computecanada.ca). *This publication is part of the Human Cell Atlas –* www.humancellatlas.org/publications.

## Data availability

The PMBC dataset is available from the 10x genomics repository (corresponding to healthy donor A) from the link https://support.10xgenomics.com/single-cell-gene-expression/datasets/1.1.0/fresh_68k_pbmc_donor_a?. The BoneMarrow dataset is available in the Gene Expression Omnibus repository under accession number <u>GSE72857</u>. A collection of simulated datasets with known ground truth GRNs will be publicly provided upon publication to enable GRN inference benchmarking on various datasets.

## Code availability

Our implementation and evaluation of GRouNdGAN in Python 3.9.6 using the PyTorch framework^60^ along with a tutorial is freely available under the GNU Affero General Public License v3.0 on GitHub (https://github.com/Emad-COMBINE-lab/GRouNdGAN). The repository also contains our PyTorch implementation of scGAN, projection-based conditioning cscGAN^24^ and a variation of cscGAN that uses conditioning by concatenation (cWGAN).

## Supplementary Material

File S1: Contains supplementary notes, all supplementary figures, Tables S1-S3, and caption of Tables S4-S6 (which are provided as separate files).

## References

1 Lee, T. I. et al. Transcriptional regulatory networks in Saccharomyces cerevisiae. science 298, 799–804 (2002).

2 Che, D. et al. Dynamic and modular gene regulatory networks drive the development of gametogenesis. Briefings in bioinformatics 18, 712–721 (2017).

3 Olson, E. N. Gene regulatory networks in the evolution and development of the heart. Science 313, 1922–1927 (2006).

4 Dibaeinia, P. & Sinha, S. SERGIO: a single-cell expression simulator guided by gene regulatory networks. Cell systems 11, 252–271. e211 (2020).

5 Yang, Y. et al. Gene knockout inference with variational graph autoencoder learning single-cell gene regulatory networks. Nucleic Acids Research, gkad450 (2023).

6 Madhamshettiwar, P. B., Maetschke, S. R., Davis, M. J., Reverter, A. & Ragan, M. A. Gene regulatory network inference: evaluation and application to ovarian cancer allows the prioritization of drug targets. Genome medicine 4, 1–16 (2012).

7 Manuel, A. M., Dai, Y., Jia, P., Freeman, L. A. & Zhao, Z. A gene regulatory network approach harmonizes genetic and epigenetic signals and reveals repurposable drug candidates for multiple sclerosis. Human Molecular Genetics 32, 998–1009 (2023).

8 Moerman, T. et al. GRNBoost2 and Arboreto: efficient and scalable inference of gene regulatory networks. Bioinformatics 35, 2159–2161 (2019).

9 Huynh-Thu, V. A., Irrthum, A., Wehenkel, L. & Geurts, P. Inferring regulatory networks from expression data using tree-based methods. PloS one 5, e12776 (2010).

10 Chan, T. E., Stumpf, M. P. & Babtie, A. C. Gene regulatory network inference from single-cell data using multivariate information measures. Cell systems 5, 251–267. e253 (2017).

11 Emad, A. & Sinha, S. Inference of phenotype-relevant transcriptional regulatory networks elucidates cancer type-specific regulatory mechanisms in a pan-cancer study. NPJ systems biology and applications 7, 9 (2021).

12 Specht, A. T. & Li, J. LEAP: constructing gene co-expression networks for single-cell RNA-sequencing data using pseudotime ordering. Bioinformatics 33, 764–766 (2017).

13 Matsumoto, H. et al. SCODE: an efficient regulatory network inference algorithm from single-cell RNA-Seq during differentiation. Bioinformatics 33, 2314–2321 (2017).

14 Papili Gao, N., Ud-Dean, S. M., Gandrillon, O. & Gunawan, R. SINCERITIES: inferring gene regulatory networks from time-stamped single cell transcriptional expression profiles. Bioinformatics 34, 258–266 (2018).

15 Aibar, S. et al. SCENIC: single-cell regulatory network inference and clustering. Nature methods 14, 1083–1086 (2017).

16 Bravo González-Blas, C., et al. SCENIC+: single-cell multiomic inference of enhancers and gene regulatory networks. Nature Methods, 1–13 (2023).

17 Kartha, V. K. et al. Functional inference of gene regulation using single-cell multi-omics. Cell genomics 2 (2022).

18 Badia-i-Mompel, P. et al. Gene regulatory network inference in the era of single-cell multi-omics. Nature Reviews Genetics, 1–16 (2023).

19 Pratapa, A., Jalihal, A. P., Law, J. N., Bharadwaj, A. & Murali, T. Benchmarking algorithms for gene regulatory network inference from single-cell transcriptomic data. Nature methods 17, 147–154 (2020).

20 Emmert-Streib, F., Dehmer, M. & Haibe-Kains, B. Gene regulatory networks and their applications: understanding biological and medical problems in terms of networks. Frontiers in cell and developmental biology 2, 38 (2014).

21 Kolmykov, S. et al. GTRD: an integrated view of transcription regulation. Nucleic acids research 49, D104–D111 (2021).

22 Xu, H. et al. ESCAPE: database for integrating high-content published data collected from human and mouse embryonic stem cells. Database 2013, bat045 (2013).

23 Kanehisa, M. & Goto, S. KEGG: kyoto encyclopedia of genes and genomes. Nucleic acids research 28, 27–30 (2000).

24 Marouf, M. et al. Realistic in silico generation and augmentation of single-cell RNA-seq data using generative adversarial networks. Nature communications 11, 1–12 (2020).

25 Sun, T., Song, D., Li, W. V. & Li, J. J. scDesign2: a transparent simulator that generates high-fidelity single-cell gene expression count data with gene correlations captured. Genome biology 22, 1–37 (2021).

26 Baruzzo, G., Patuzzi, I. & Di Camillo, B. SPARSim single cell: a count data simulator for scRNA-seq data. Bioinformatics 36, 1468–1475 (2020).

27 Schaffter, T., Marbach, D. & Floreano, D. GeneNetWeaver: in silico benchmark generation and performance profiling of network inference methods. Bioinformatics 27, 2263–2270 (2011).

28 Marbach, D. et al. Revealing strengths and weaknesses of methods for gene network inference. Proceedings of the national academy of sciences 107, 6286–6291 (2010).

29 Marbach, D. et al. Wisdom of crowds for robust gene network inference. Nature methods 9, 796–804 (2012).

30 Chen, S. & Mar, J. C. Evaluating methods of inferring gene regulatory networks highlights their lack of performance for single cell gene expression data. BMC bioinformatics 19, 1–21 (2018).

31 Kocaoglu, M., Snyder, C., Dimakis, A. G. & Vishwanath, S. Causalgan: Learning causal implicit generative models with adversarial training. *arXiv preprint arXiv:1709.02023* (2017).

32 Arjovsky, M., Chintala, S. & Bottou, L. in International conference on machine learning. 214-223 (PMLR).

33 Zheng, G. X. et al. Massively parallel digital transcriptional profiling of single cells. Nature communications 8, 1–12 (2017).

34 Paul, F. et al. Transcriptional heterogeneity and lineage commitment in myeloid progenitors. Cell 163, 1663–1677 (2015).

35 Gretton, A., Borgwardt, K. M., Rasch, M. J., Schölkopf, B. & Smola, A. A kernel two-sample test. The Journal of Machine Learning Research 13, 723–773 (2012).

36 Korsunsky, I. et al. Fast, sensitive and accurate integration of single-cell data with Harmony. Nature methods 16, 1289–1296 (2019).

37 Kim, S. ppcor: an R package for a fast calculation to semi-partial correlation coefficients. Communications for statistical applications and methods 22, 665 (2015).

38 Wolf, F. A. et al. PAGA: graph abstraction reconciles clustering with trajectory inference through a topology preserving map of single cells. Genome biology 20, 1–9 (2019).

39 Haghverdi, L., Büttner, M., Wolf, F. A., Buettner, F. & Theis, F. J. Diffusion pseudotime robustly reconstructs lineage branching. Nature methods 13, 845–848 (2016).

40 Trapnell, C. et al. Pseudo-temporal ordering of individual cells reveals dynamics and regulators of cell fate decisions. Nature biotechnology 32, 381 (2014).

41 Kumar, P., Tan, Y. & Cahan, P. Understanding development and stem cells using single cell-based analyses of gene expression. Development 144, 17–32 (2017).

42 Saelens, W., Cannoodt, R., Todorov, H. & Saeys, Y. A comparison of single-cell trajectory inference methods. Nature biotechnology 37, 547–554 (2019).

43 Street, K. et al. Slingshot: cell lineage and pseudotime inference for single-cell transcriptomics. BMC genomics 19, 1–16 (2018).

44 Hwang, B., Lee, J. H. & Bang, D. Single-cell RNA sequencing technologies and bioinformatics pipelines. Experimental & molecular medicine 50, 1–14 (2018).

45 Jacomy, M., Venturini, T., Heymann, S. & Bastian, M. ForceAtlas2, a continuous graph layout algorithm for handy network visualization designed for the Gephi software. PloS one 9, e98679 (2014).

46 Wolf, F. A., Angerer, P. & Theis, F. J. SCANPY: large-scale single-cell gene expression data analysis. Genome biology 19, 1–5 (2018).

47 Gao, Y., Shen, L. & Xia, S.-T. in ICASSP 2021-2021 IEEE International Conference on Acoustics, Speech and Signal Processing (ICASSP). 3320–3324 (IEEE).

48 Satija, R., Farrell, J. A., Gennert, D., Schier, A. F. & Regev, A. Spatial reconstruction of single-cell gene expression data. Nature biotechnology 33, 495–502 (2015).

49 Goodfellow, I. et al. Generative adversarial networks. Communications of the ACM 63, 139–144 (2020).

50 Gulrajani, I., Ahmed, F., Arjovsky, M., Dumoulin, V. & Courville, A. C. Improved training of wasserstein gans. Advances in neural information processing systems 30 (2017).

51. Reddi, S. J., Kale, S. & Kumar, S. On the convergence of adam and beyond. *arXiv preprint arXiv:1904.09237* (2019).

52 He, K., Zhang, X., Ren, S. & Sun, J. in Proceedings of the IEEE international conference on computer vision. 1026–1034.

53 Glorot, X. & Bengio, Y. in Proceedings of the thirteenth international conference on artificial intelligence and statistics. 249-256 (JMLR Workshop and Conference Proceedings).

54 Ioffe, S. & Szegedy, C. in International conference on machine learning. 448-456 (PMLR).

55 Hu, H. et al. AnimalTFDB 3.0: a comprehensive resource for annotation and prediction of animal transcription factors. Nucleic acids research 47, D33–D38 (2019).

56 Shaham, U. et al. Removal of batch effects using distribution-matching residual networks. Bioinformatics 33, 2539–2546 (2017).

57 Shu, H. et al. Modeling gene regulatory networks using neural network architectures. Nature Computational Science 1, 491–501 (2021).

58 Mirza, M. & Osindero, S. Conditional generative adversarial nets. *arXiv preprint arXiv:1411*.1784 (2014).

59 Islam, S. et al. Characterization of the single-cell transcriptional landscape by highly multiplex RNA-seq. Genome research 21, 1160–1167 (2011).

60 Paszke, A. et al. Pytorch: An imperative style, high-performance deep learning library. Advances in neural information processing systems 32 (2019).

