## Supplementary material for "GRouNdGAN: GRN-guided simulation of single-cell RNA-seq data using causal generative adversarial networks": File S1

3480 University Street, Montreal, Quebec, Canada, H3A 0E9

### Supplementary notes

#### An ablation study for different components of GRouNdGAN

GRouNdGAN includes several components and incorporates auxiliary networks to provide a realistic simulated dataset and to ensure that the causal GRN is imposed within the generated cells. Here, we investigated the effect that these components (labeler, anti-labeler, library size normalization layer, and noise input to the target generators) have on the causal GRN enforcement and on the quality of data simulation. We conducted this ablation study on the BoneMarrow dataset. For the imposed GRN, we first identified top ten TFs for each gene using GRNBoost2 (using the real dataset), but only imposed the regulatory edge between the gene and its top 1$^{st}$, 3$^{rd}$, 5$^{th}$, 7$^{th}$, and 9$^{th}$ TFs. The small sample size of this dataset and the imposition of a GRN that to some degree deviates from the GRN that best matches the real reference single-cell dataset make training a model on this (dataset, GRN) combination more challenging, allowing us to observe more granularity when ablating from our model. Supplementary Table S3 shows the effect of each change in the architecture of the model on the performance. It is evident from this table that GRouNdGAN performs best across all metrics with a labeler, anti-labeler, and target generator input noises. Not performing the library-size normalization on the input dataset and removing the LSN layer results in higher miLISI and lower Euclidean distance, but the MMD, Cosine distance, AUPRC of GRN inference, and the AUROC of RF all deteriorate. Additionally, including these steps in the model results in smoother training.

#### The stability of GRouNdGAN and the effect of GRN properties on its performance

To assess the stability of GRouNdGAN, we generated five different batches (or “replicates”) of 1000 cells using the PBMC-CTL dataset. Comparing each generated replicate with the held-out real test set (Supplementary Table S5) reveals a stable performance across different generated replicates and a high degree of resemblance to real data.

Next, we sought to determine the effect of the choice of the imposed GRN on the simulated data. We imposed five different GRNs with different properties, and simulated data using the PBMC-CTL and the BoneMarrow datasets (Supplementary Table S5). Comparing the performance metrics measuring resemblance of the simulated and real samples, we observed that reducing the number of TFs regulating each gene results in some deterioration of the performance. We hypothesized that this is because when we reduce the number of regulating TFs, the GRN is less representative of the TF-gene relationships of the real dataset. To test this (while controlling for the number of regulating TFs), we compared the dataset generated by a GRN composed of the top ten TFs for each gene and the dataset generated by a GRN composed of the bottom ten TFs for each gene (edges that were ranked very low by GRNBoost2 on the experimental dataset). Consistent with our hypothesis, the former dataset had much better performance (Supplementary Table S5). For example, data generated based on PBMC-CTL and top 10 TFs had an miLIS = 1.89 and MMD=0.026 (on test set), while the data generated based on bottom 10 TFs an miLIS = 1.56 and MMD=0.355.

The reason for this behavior is that when generating realistic simulated data, imposing a GRN that does not conform to the underlying TF-gene relationship in the real dataset makes the task extremely challenging. This is because incorporating the causal GRN imposes certain patterns in the gene expression profiles of samples. If this pattern is drastically different from the one present in the real dataset, generating realistic samples and imposing the GRN simultaneously act as contradictory requirements. This is the reason that simulators that do not impose a GRN in the data, have a much easier task, as they only need to generate realistic synthetic data even if they disrupt the TF-gene (or co-expression) relationships. In fact, we observed that when a GRN inference method (GRNBoost2) was applied to data generated by scGAN, its performance was much lower than when they were applied to real data (Figure 3), showing that some TF-gene relationships are disrupted by the simulator. In spite of this deterioration in performance, when a reasonable number of top TFs were selected (e.g., top 5 TFs), a good performance could be achieved (e.g., the test set miLISI was larger than 1.80 for both datasets).

**Details regarding the Wasserstein distance**

Various metrics exist to quantify the similarity between the generator's and real data's probability distributions. While initial implementations of the GAN used Kullback–Leibler or Jensen–Shannon divergence, they are prone to model collapse where the generator learns to produce only a subset of modes in the dataset[1-4]. Numerous approaches have been proposed to solve this problem, from regularization technique to using separate generators, explicitly enforcing the GAN to learn all modes[1,4-6], etc. Wasserstein distance, when used as the divergence metric in GANs has led to stability in training without evidence of mode collapse[7]. Mathematically, the Wasserstein or Earth-Mover distance is defined as

$$W(\mathbb{P}_r, \mathbb{P}_g) = \inf_{\gamma \in \Pi(\mathbb{P}_r, \mathbb{P}_g)} \mathbb{E}_{(x,y)\sim\gamma}[\, \|x - y\| \,],$$

where $\Pi(\mathbb{P}_r, \mathbb{P}_g)$ is the set of all joint distributions over $x$ and $y$ whose marginals are respectively $\mathbb{P}_r$ and $\mathbb{P}_g$. Intuitively, $\gamma(x, y)$ denotes the unit of mass to be transported from $x$ to $y$ to transform $\mathbb{P}_r$ to $\mathbb{P}_g$ representing the optimal transport plan.

However, the infimum in equation 1 is intractable over all $\gamma \in \Pi(\mathbb{P}_\mathrm{r}, \mathbb{P}_\mathrm{g})$, for this reason an equivalent formulation of the Wasserstein distance from the Kantorovich-Rubinstein duality[8] can be obtained,

$$W(\mathbb{P}_r, \mathbb{P}_g) = \sup_{\|f\|_L \leq 1} \mathbb{E}_{x\sim\mathbb{P}_r}[f(x)] - \mathbb{E}_{x\sim\mathbb{P}_g}[f(x)],$$

where $W(\mathbb{P}_r, \mathbb{P}_g)$ is the supremum over all 1-Lipschitz functions with values in $\mathbb{R}$ ($f: X \rightarrow \mathbb{R}$). To enforce a Lipschitz constraint on the critic, we added a gradient penalty term proposed by Gulrajani et al.[2] as an alternative to weight clipping used in the original WGAN which is shown to suffer from vanishing or exploding gradients and capacity underuse. We approximated the solution of this network by the discriminative network (critic).

**Supplementary tables**

**Table S1:** Architectural choices and hyperparameters of the WGAN used to pre-train the causal controller (Figure 1A).

| **Hyperparameter / Architectural Choice** | **Value** |
|---|---|
| Length of the input noise vector (latent dimension) | 128 |
| Width of generator's input layer | 128 |
| Width of generator's hidden layers | [256, 512, 1024] |
| Width of generator's output layer | 1000 |
| Width of critic's input layer | 1000 |
| Width of critic's hidden layers | [1024, 512, 256] |
| Width of critic's output layer | 1 |
| Library size | 20000 |
| Regularization parameter of the gradient penalty ($\lambda$) | 10 |
| Batch size | 128 |
| Number of critic iterations per generator one iteration | 5 |
| Number of training steps | 200000 |
| AMSGrad's coefficients for computing the running averages of the gradient and its square | $\beta_1 = 0.5, \beta_2 = 0.9$ |
| Generator's initial learning rate | 0.0001 |
| Generator's final learning rate | 0.00001 |
| Critic's initial learning rate | 0.0001 |
| Critic's final learning rate | 0.00001 |

**Table S2:** Architectural choices and hyperparameters used in GRouNdGAN (Figure 1B).

| **Hyperparameter / Architectural Choice** | **Value** |
|---|---|
| Dimension of the noise variable per target generator ($N_{noise}$) | 1 |
| Width of the input layer for the generator of a target gene with $N_{TF}$ regulating TFs | $N_{noise} + N_{TF}$ |
| Number of hidden layers of each target generator | 3 |
| Width multiplier ($w$) | 2 |
| Width of each target generator's hidden layer | $w\ (N_{noise} + N_{TF})$ |
| Size of the output layer for each target generator | 1 |
| Width of the critic's input layer | 1000 |
| Width of critic's hidden layers | [1024, 512, 256] |
| Size of the critic's output layer | 1 |
| Width of the input layer for the Labeler and the Anti-Labeler | Number of target genes |
| Width of the hidden layers of the Labeler and the Anti-Labeler | [2000, 2000, 2000] |
| Width of the Labeler and the Anti-Labeler output layer | Total Number of TFs |
| Library size | 20000 |
| Regularization parameter of the gradient penalty ($\lambda$) | 10 |
| Batch size | 1024 |
| Number of critic iterations per generator one iteration | 5 |
| Number of training steps | 700000 |
| AMSGrad's coefficients for computing the running averages of the gradient and its square | $\beta_1 = 0.5$, $\beta_2 = 0.9$ |
| Generator's initial learning rate | 0.001 |
| Generator's final learning rate | 0.0001 |
| Critic's initial learning rate | 0.001 |
| Critic's final learning rate | 0.0001 |

**Table S3:** The results of the ablation study on a held-out test set of the BoneMarrow dataset. The first column shows the variation of the model. The values in the table show the percentage of the change in the performance metrics compared to GrouNdGAN. The first five metrics quantify the resemblance of simulated and experimental data. The last column shows the decrease in the AUPRC of GRN inference using simulated data (using GRNBoost2) and quantifies the effect of ablation on GRN imposition.

| **Model** | **Increase in RF AUROC (%)** | **Increase in Cosine distance (%)** | **Increase in Euclidean distance (%)** | **Decrease in miLISI (%)** | **Increase in MMD (%)** | **Decrease in GRN inference AUPRC (%)** |
|---|---|---|---|---|---|---|
| GrouNdGAN w/o labeler | 16.7 | 4.5 | 7.5 | 40.5 | 16.7 | 5.2 |
| GrouNdGAN w/o anti-labeler | 50.0 | 16.7 | 4.0 | 39.3 | 50.0 | 6.5 |
| GrouNdGAN w/o labeler & w/o anti-labeler | 7.6 | 100.0 | 54.9 | 1.5 | 61.9 | 7.8 |
| GrouNdGAN w/o noise to target generators | 16.5 | 33.3 | 25.4 | 7.7 | 54.8 | 2.6 |
| GrouNdGAN w/o LSN layer & w/o input normalization | 21.5 | 50.0 | -67.6 | -3.3 | 23.8 | 5.2 |

**Table S4:** The performance of different simulators in generating realistic simulated data using three datasets. The table is provided as a separate xlsx file.

**Table S5:** Stability analysis and the effect of different GRN properties on the performance. The table is provided as a separate xlsx file.

**Table S6:** Results of GRN inference using different datasets simulated by GrouNdGAN. The table is provided as a separate xlsx file.

**Table S7:** Cell types present in the BoneMarrow dataset, their abbreviations, and the list of marker genes associated with each used to annotate nodes of the PAGA graphs. The markers genes were obtained from the link (https://scanpy-tutorials.readthedocs.io/en/latest/paga-paul15.html) [9]. Marker genes not included in the top 1000 highly variable genes were excluded from this table and not used in our analyses.

| **Cell type** | **Abbreviation** | **Marker gene(s)** |
|---|---|---|
| Erythroid cells | Ery | *Gata1, Klf1, Hba-a2* |
| Neutrophils | Neu | *Elane, Cebpe, Ctsg, Mpo* |
| Monocytes | Mo | *Irf8, Csf1r, Ctsg, Mpo* |
| Megakaryocytes | Mk | *Itga2b* (encodes protein CD41), *Pbx1, Sdpr, Vwf* |
| Basophils | Baso | *Mcpt8, Prss34* |
| Mast cells | Ma | *Cma1, Gzmb* |
| Mast cells & Basophils | Ma_Baso | *Cpa3* |

**Table S8:** The effect of cell type specific TF knockout on the miLISI values of each cell type. The first column shows the cell type, the second column shows the percentage of that cell type in the data, the third and fourth columns show miLISI before and after knockout (KO) of top three TFs (shown in the seventh column), respectively. The fifth and sixth columns show the miLISI of other cell types before and after the KO experiment.

| **Cell type** | **Perc.** | **miLISI before KO** | **miLISI after KO** | **miLISI of other cells before KO** | **miLISI of other cells after KO** | **Top DE TFs** |
|---|---|---|---|---|---|---|
| CD14+ Monocytes | 4.75 | 1.97 | 1.42 | 1.94 | 1.91 | SPI1, CEBPD, JUND |
| CD19+ B cells | 8.7 | 1.96 | 1.42 | 1.94 | 1.92 | SPIB, IRF8, LYL1 |
| CD56+ Natural Killer (NK) cells | 11.8 | 1.93 | 1.71 | 1.95 | 1.92 | HOPX, ASCL2, ZNF683 |
| Dendritic cells | 3.4 | 1.98 | 1.46 | 1.94 | 1.89 | SPI1, CEBPD, LYL1 |

**Supplementary figures**

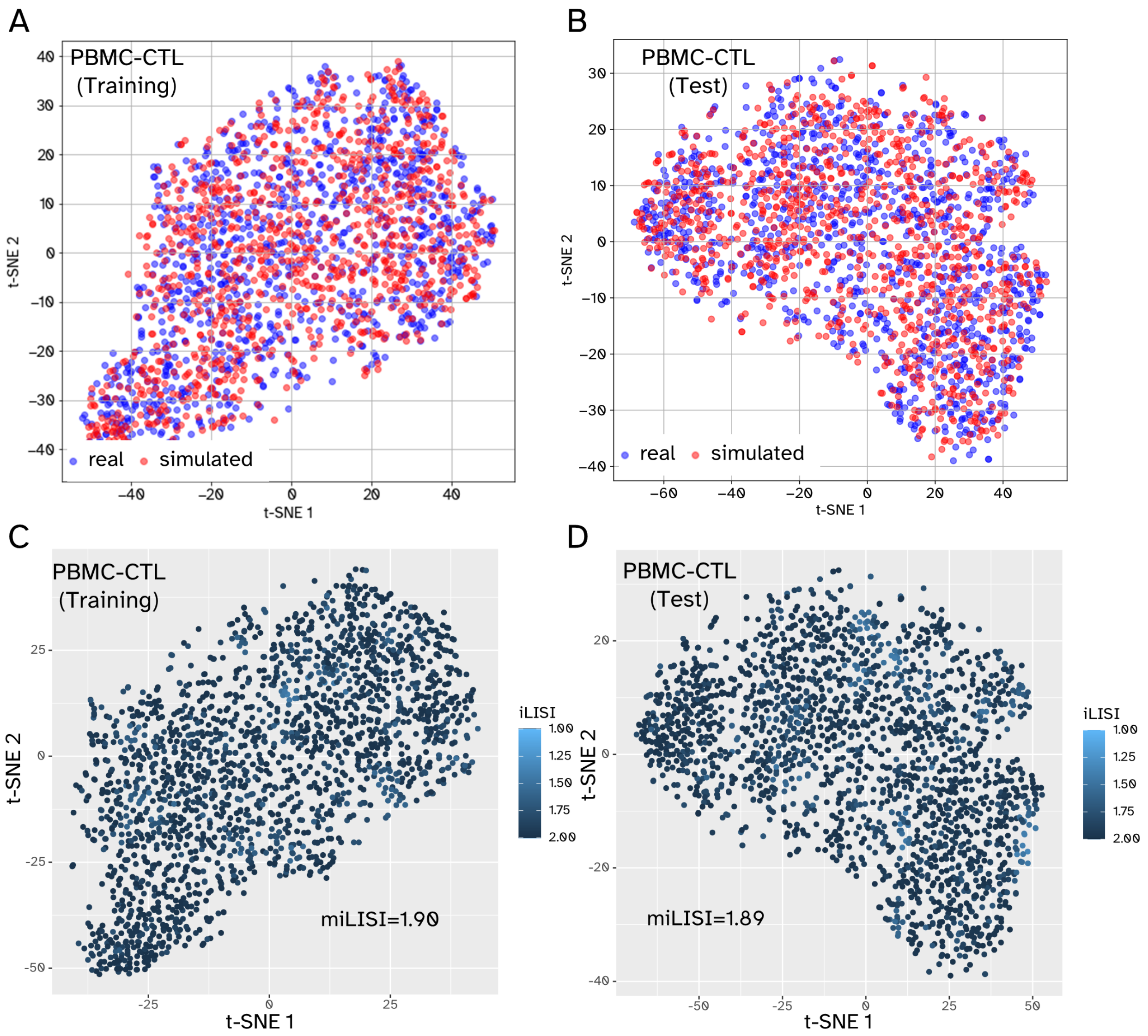


**Figure S1:** Real (experimental) and GRouNdGAN-simulated scRNA-seq data based on the PBMC-CTL dataset. All plots correspond to 1000 simulated cells and 1000 real cells. Each gene in the GRN of GRouNdGAN is regulated by 15 TFs (identified using GRNBoost2 from the experimental training dataset). The top row of panels shows t-SNE plots of simulated cells (red) and real cells (blue). The bottom row of panels shows the iLISI values of each datapoint and the average iLISI score of the data (miLISI). The panels on the left correspond to comparison between simulated cells and a random set of real cells in the training set, while the panels on the right correspond to comparison between simulated cells and all the real cells in the test set.

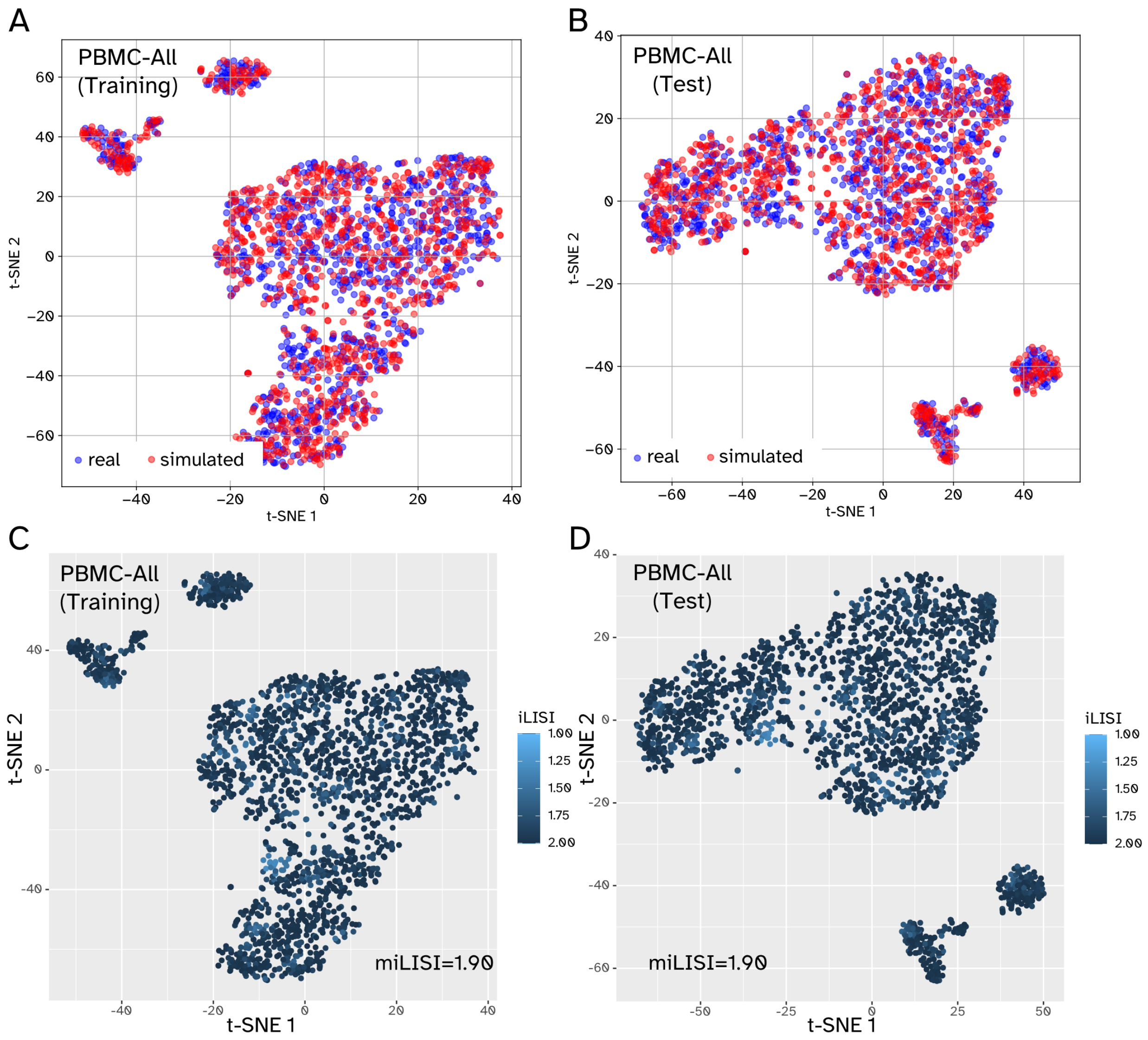


**Figure S2:** Real (experimental) and GRouNdGAN-simulated scRNA-seq data using the PBMC-All dataset. All plots correspond to 1000 simulated cells and 1000 real cells. Each gene in the GRN of GRouNdGAN is regulated by 15 TFs (identified using GRNBoost2 from the experimental training dataset). The top row of panels shows t-SNE plots of simulated cells (red) and real cells (blue). The bottom row of panels shows the iLISI values of each datapoint and the average iLISI score of the data (miLISI). The panels on the left correspond to comparison between simulated cells and a random set of real cells in the training set, while the panels on the right correspond to comparison between simulated cells and all the real cells in the test set.

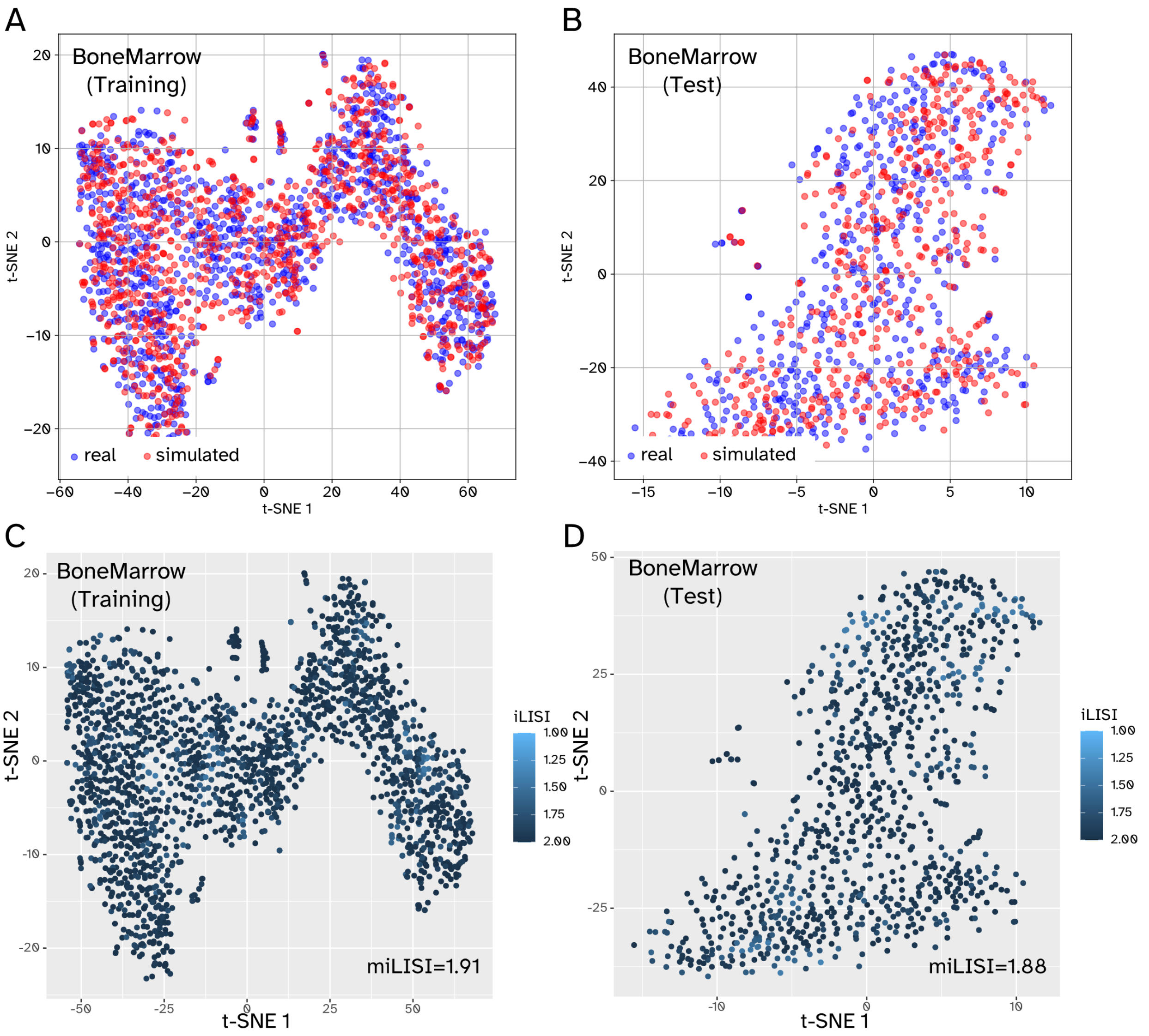


**Figure S3:** Real experimental and GRouNdGAN-simulated scRNA-seq data using the BoneMarrow dataset. All plots correspond to 500 simulated cells and 500 real cells. Each gene in the GRN of GRouNdGAN is regulated by 15 TFs (identified using GRNBoost2 from the experimental training dataset). The top row of panels shows t-SNE plots of simulated cells (red) and real cells (blue). The bottom row of panels shows the iLISI values of each datapoint and the average iLISI score of the data (miLISI). The panels on the left correspond to comparison between simulated cells and a random set of real cells in the training set, while the panels on the right correspond to comparison between simulated cells and all the real cells in the test set.

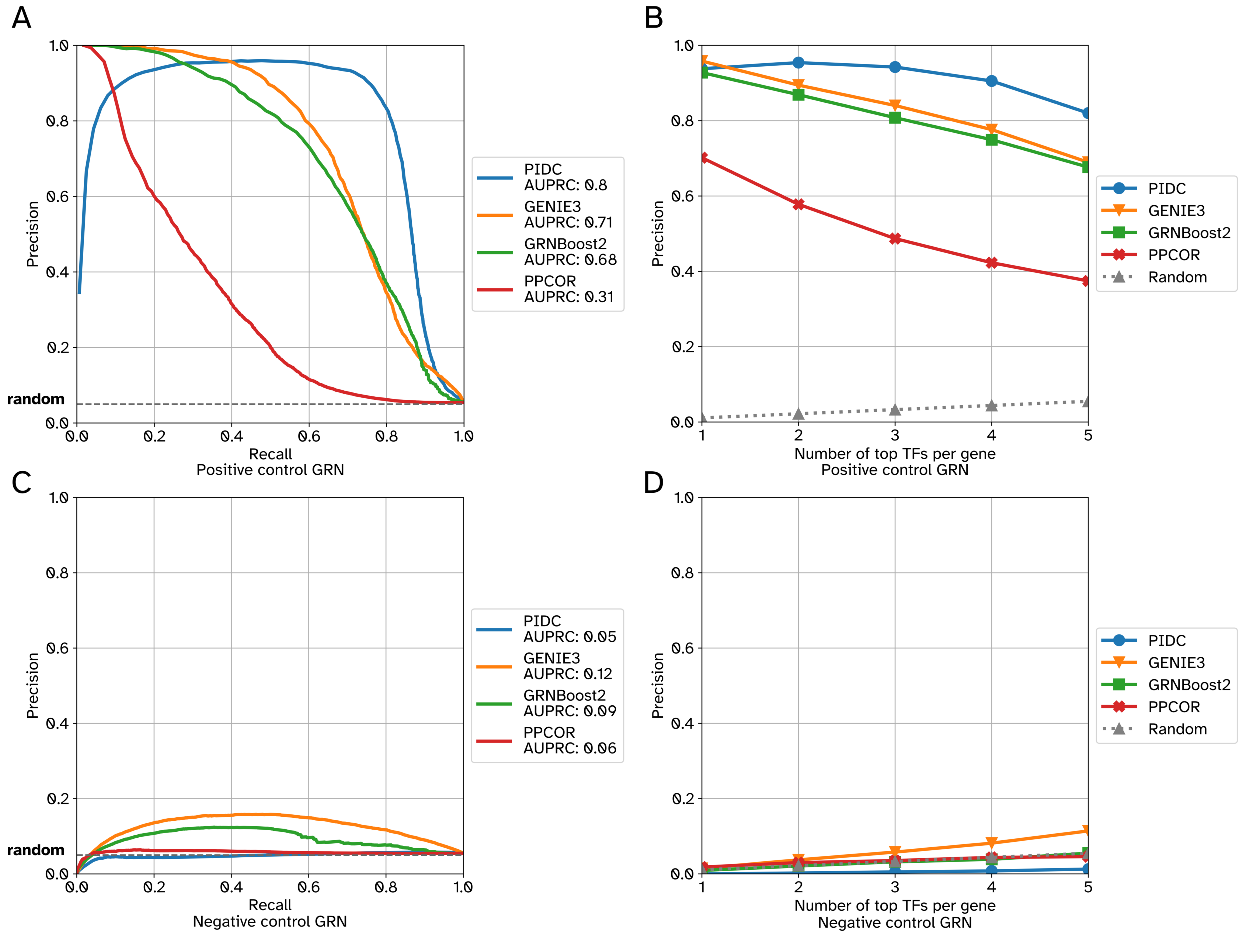


**Figure S4:** Performance of different GRN inference algorithms in recovering the imposed edges versus unimposed edges using data generated by GRouNdGAN based on the PBMC-CTL dataset. Top row shows the AUPRC (A) and Precision at k (per gene) (B) when the imposed edges (positive control GRN) were considered the ground truth. Bottom row shows the AUPRC (C) and Precision at k (per gene) (D) when the unimposed edges (negative control GRN) were considered the ground truth. Precision at k (per gene) refers to the precision when top k TFs for each gene is used to form the reconstructed GRN.

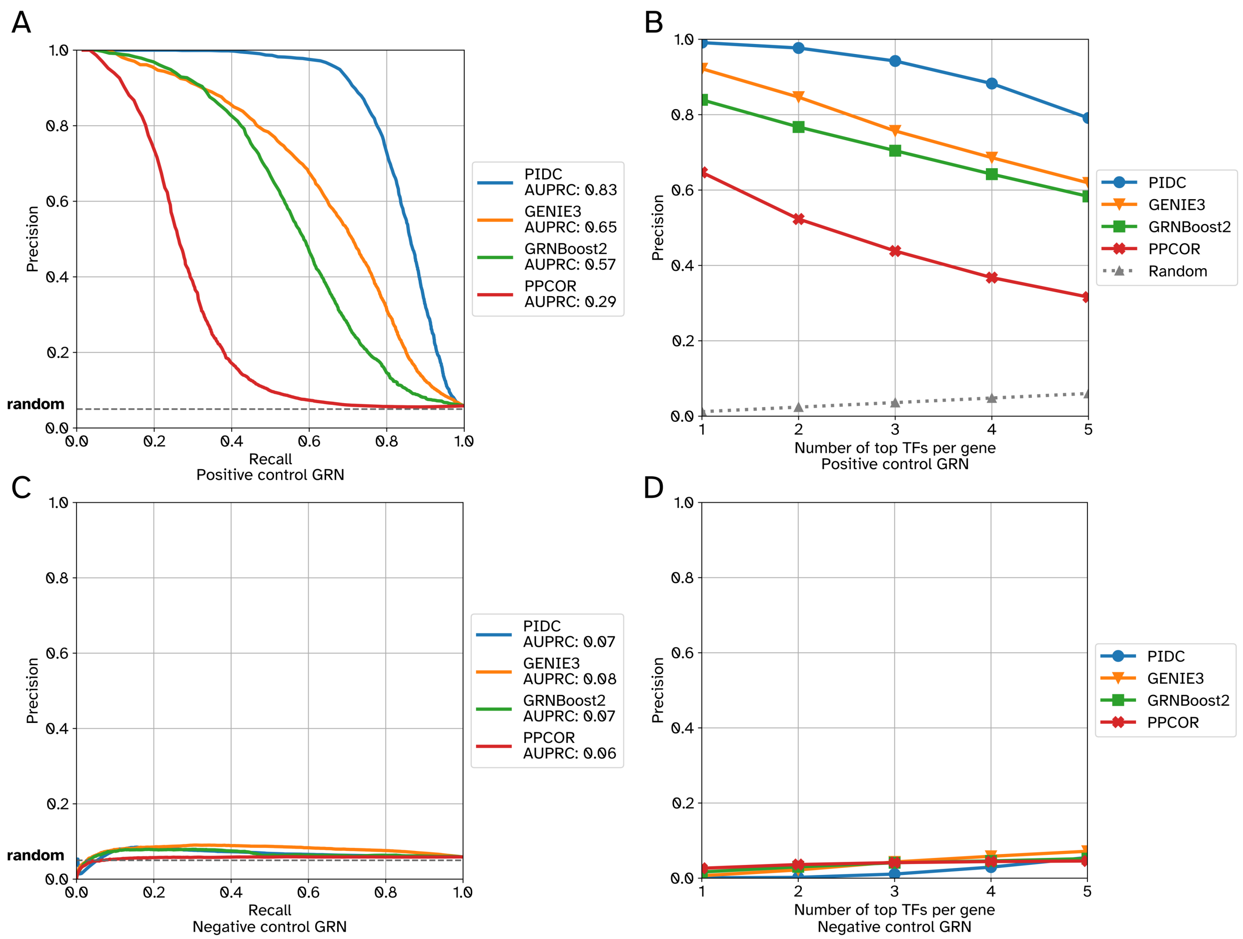


**Figure S5:** Performance of different GRN inference algorithms in recovering the imposed edges versus unimposed edges using data generated by GRouNdGAN based on the BoneMarrow dataset. Top row shows the AUPRC (A) and Precision at k (per gene) (B) when the imposed edges (positive control GRN) were considered the ground truth. Bottom row shows the AUPRC (C) and Precision at k (per gene) (D) when the unimposed edges (negative control GRN) were considered the ground truth. Precision at k (per gene) refers to the precision when top k TFs for each gene is used to form the reconstructed GRN.

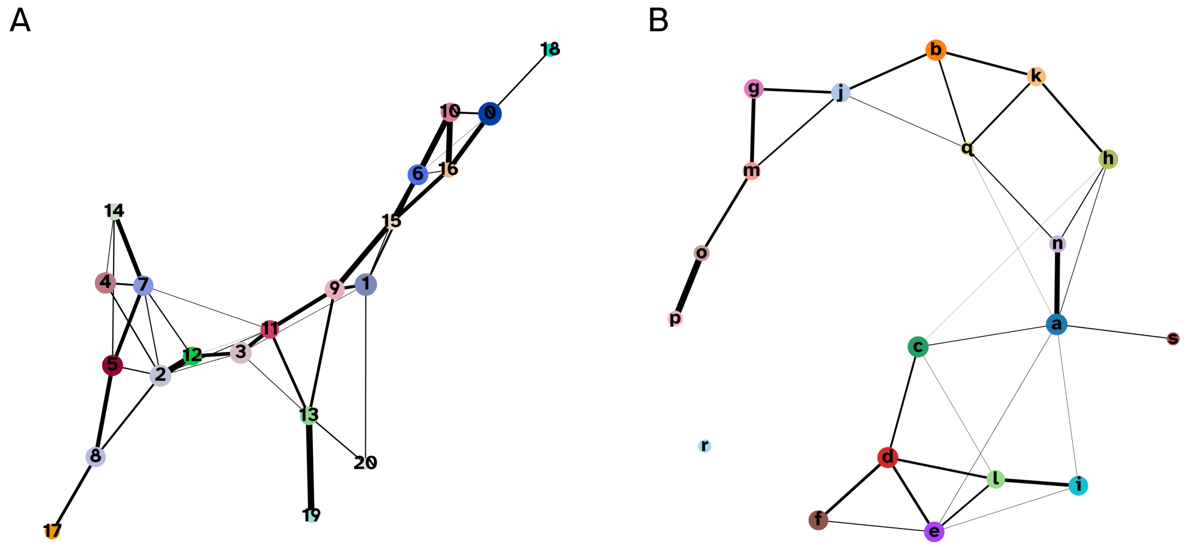


**Figure S6:** PAGA generated graphs for the BoneMarrow dataset. Nodes represent Louvain clusters capturing discrete states, edges show transitions among these states, and edge weights show-case the confidence in the existence of connections. A) The PAGA graph computed from GRouNdGAN-generated data comprising of 21 clusters. Each gene in the GRN of GRouNdGAN is regulated by 15 TFs (identified using GRNBoost2 from the real dataset) and the same number of cells as the original dataset was generated. B) The PAGA graph computed from the original BoneMarrow dataset comprising of 19 clusters. Low-connectivity edges below a threshold of 0.01 were discarded.

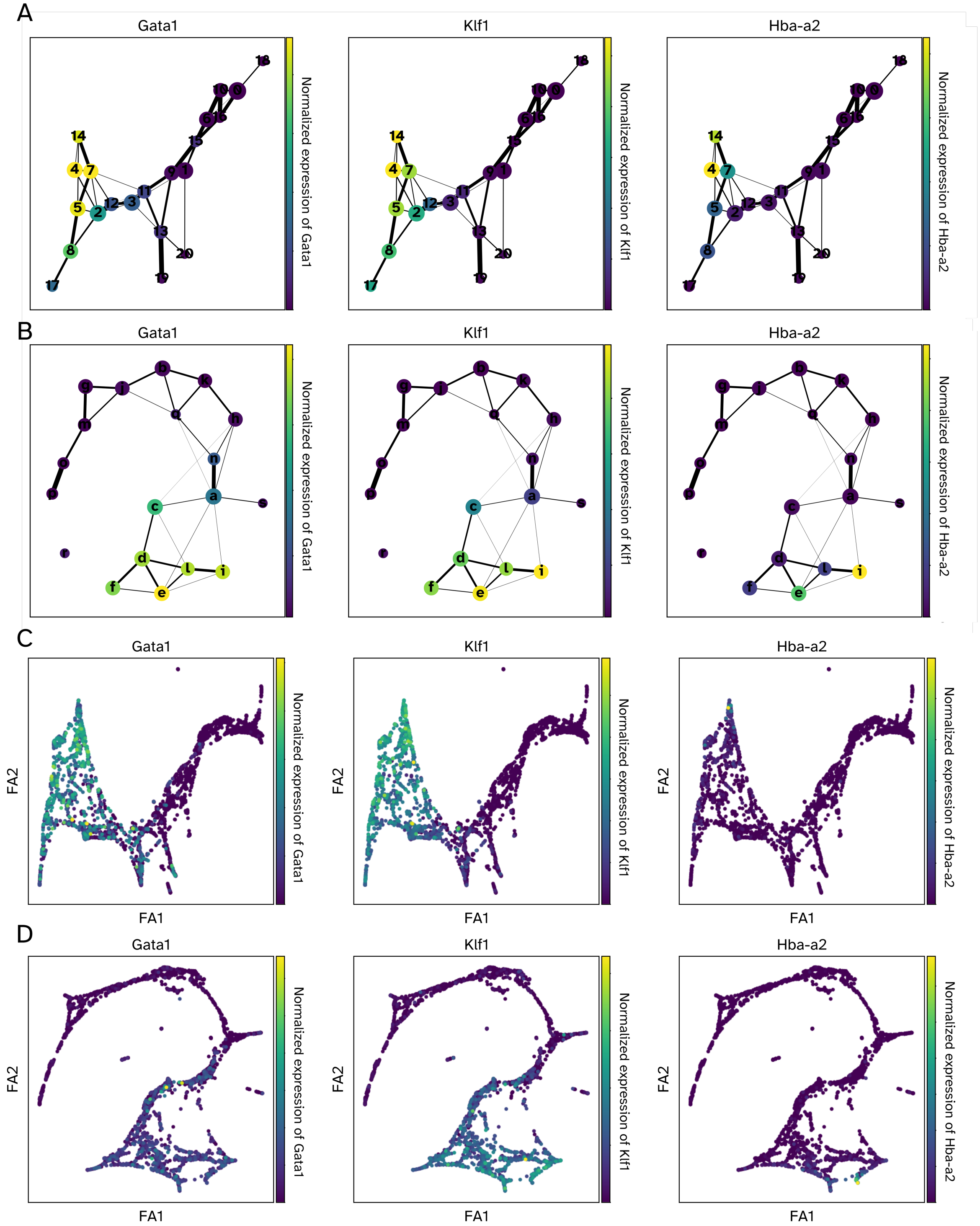


**Figure S7:** Erythroid cells' marker genes activation patterns in experimental and simulated data. Panels A and C show the normalized gene expression of the marker genes in PAGA graphs for the simulated and the experimental data, respectively. Nodes correspond to Louvain clusters capturing discrete states and edges show transitions among these states. Panels B and D show PAGA-initialized single-cell embeddings obtained using ForceAtlas2. In all figures, colors yellow and purple show the highest and lowest normalized gene expression value, respectively.

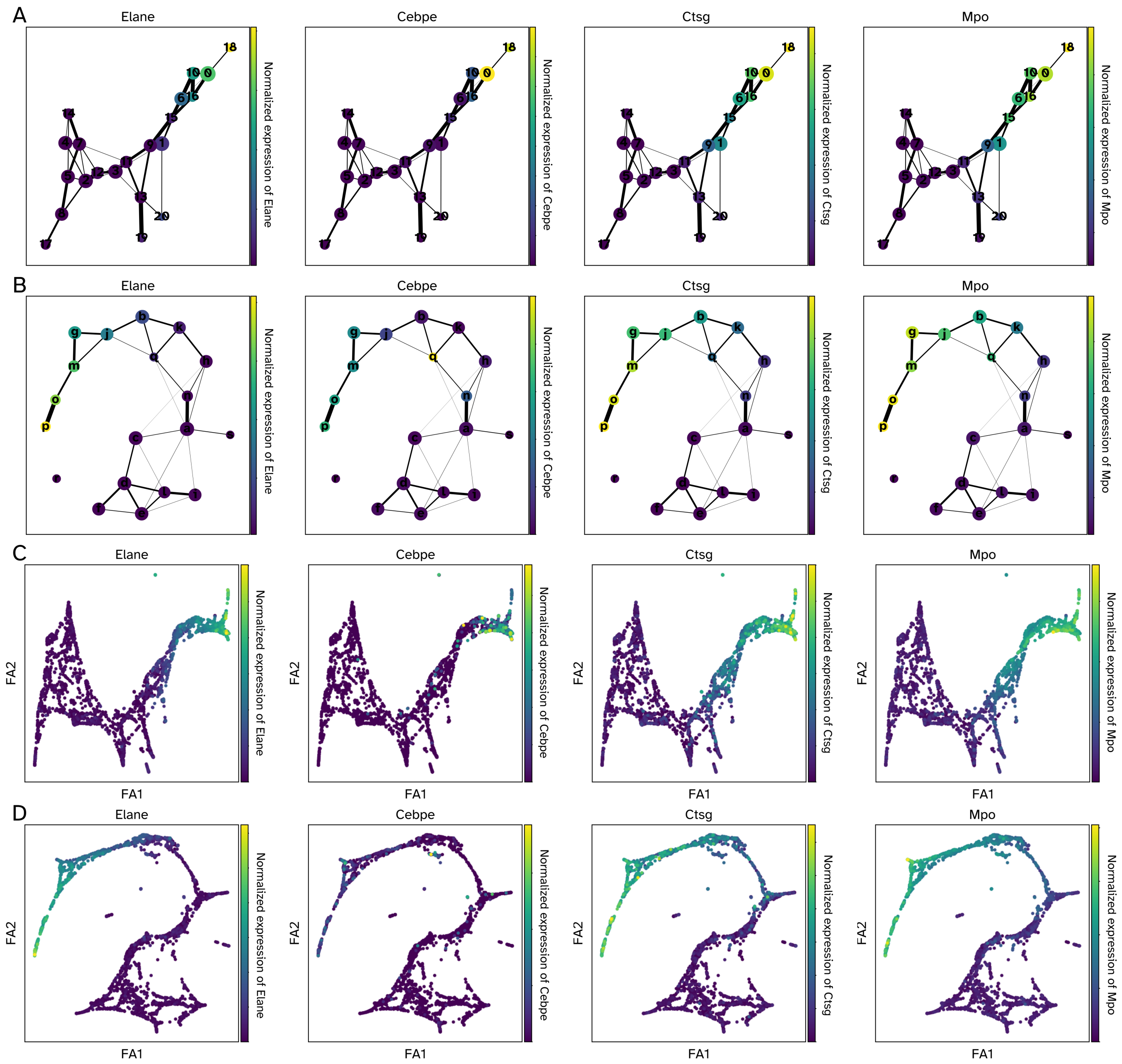


**Figure S8:** Neutrophils' marker genes activation patterns in experimental and simulated data. Panels A and C show the normalized gene expression of the marker genes in PAGA graphs for the simulated and the experimental data, respectively. Nodes correspond to Louvain clusters capturing discrete states and edges show transitions among these states. Panels B and D show PAGA-initialized single-cell embeddings obtained using ForceAtlas2. In all figures, colors yellow and purple show the highest and lowest normalized gene expression value, respectively.

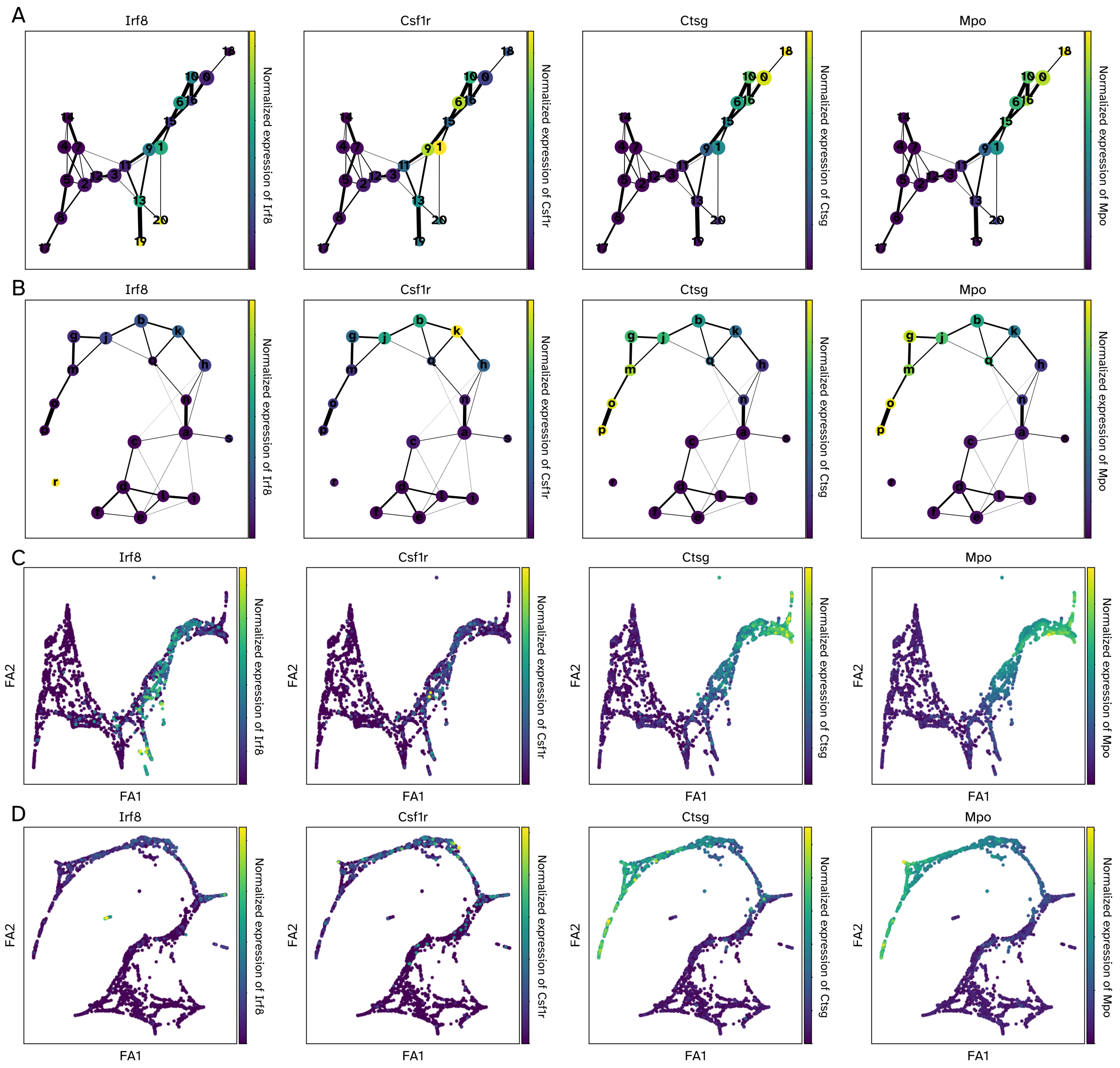


**Figure S9:** Monocytes' marker genes activation patterns in experimental and simulated data. Panels A and C show the normalized gene expression of the marker genes in PAGA graphs for the simulated and the experimental data, respectively. Nodes correspond to Louvain clusters capturing discrete states and edges show transitions among these states. Panels B and D show PAGA-initialized single-cell embeddings obtained using ForceAtlas2. In all figures, colors yellow and purple show the highest and lowest normalized gene expression value, respectively.

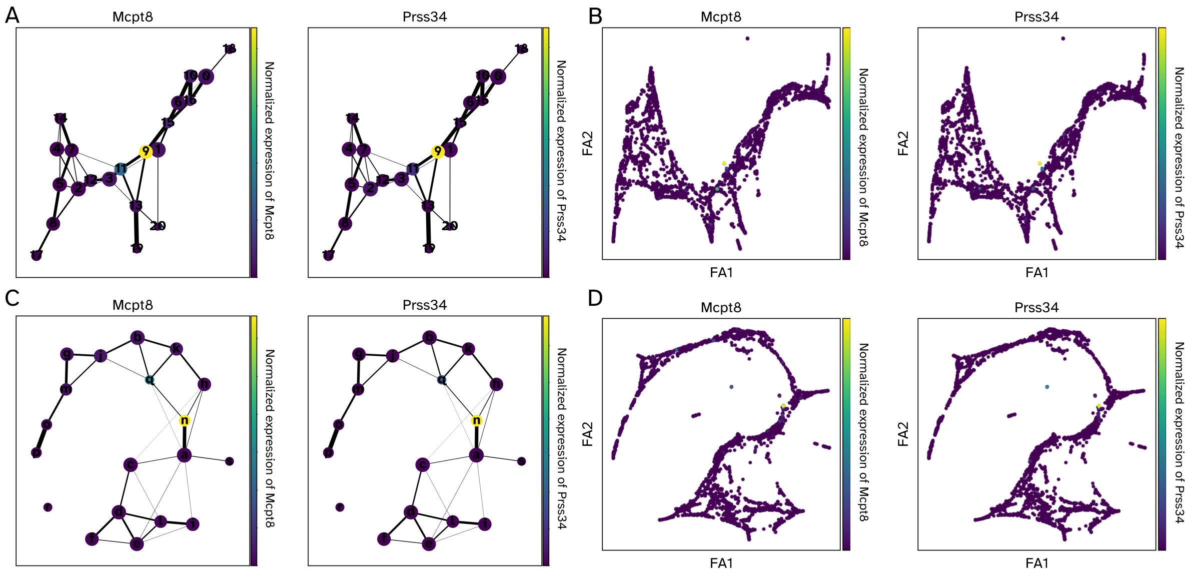


**Figure S10:** Basophils' marker genes activation patterns in experimental and simulated data. Panels A and C show the normalized gene expression of the marker genes in PAGA graphs for the simulated and the experimental data, respectively. Nodes correspond to Louvain clusters capturing discrete states and edges show transitions among these states. Panels B and D show PAGA-initialized single-cell embeddings obtained using ForceAtlas2. In all figures, colors yellow and purple show the highest and lowest normalized gene expression value, respectively.

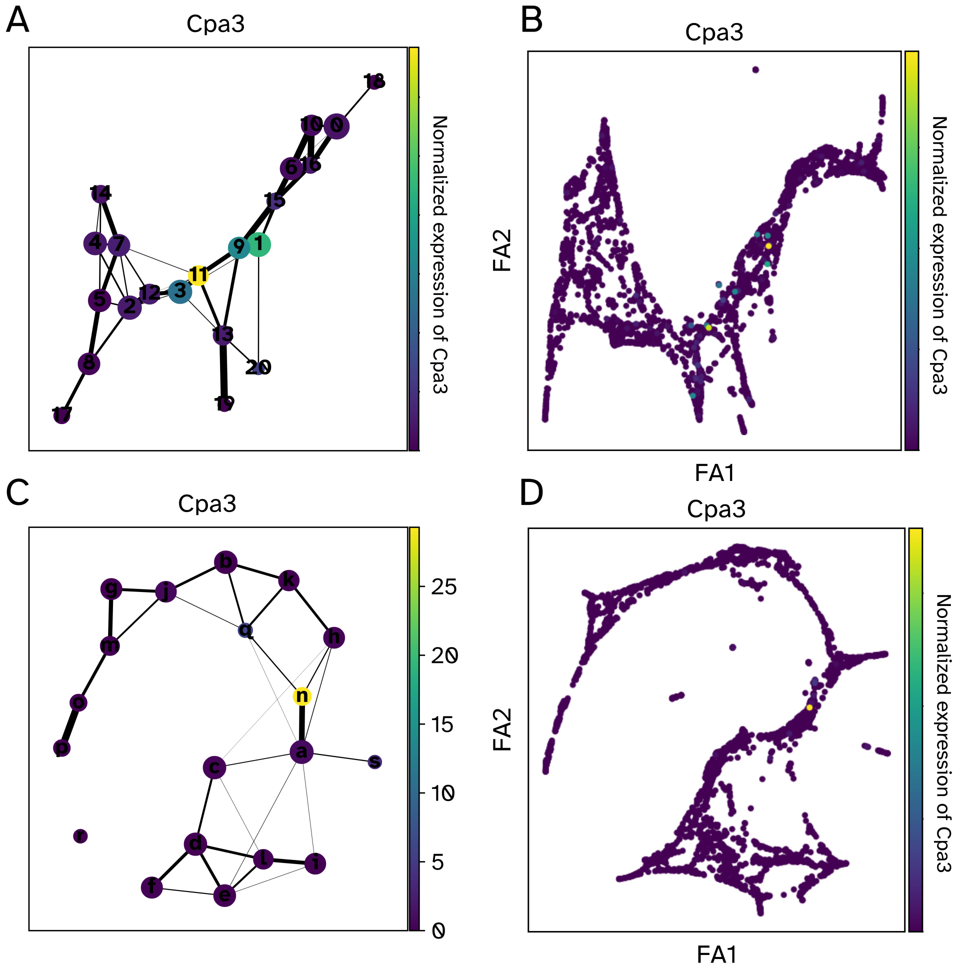


**Figure S11:** Basophils and Mast cells' marker gene activation patterns in experimental and simulated data. Panels A and C show the normalized gene expression of the marker gene in PAGA graphs for the simulated and the experimental data, respectively. Nodes correspond to Louvain clusters capturing discrete states and edges show transitions among these states. Panels B and D show PAGA-initialized single-cell embeddings obtained using ForceAtlas2. In all figures, colors yellow and purple show the highest and lowest normalized gene expression value, respectively.

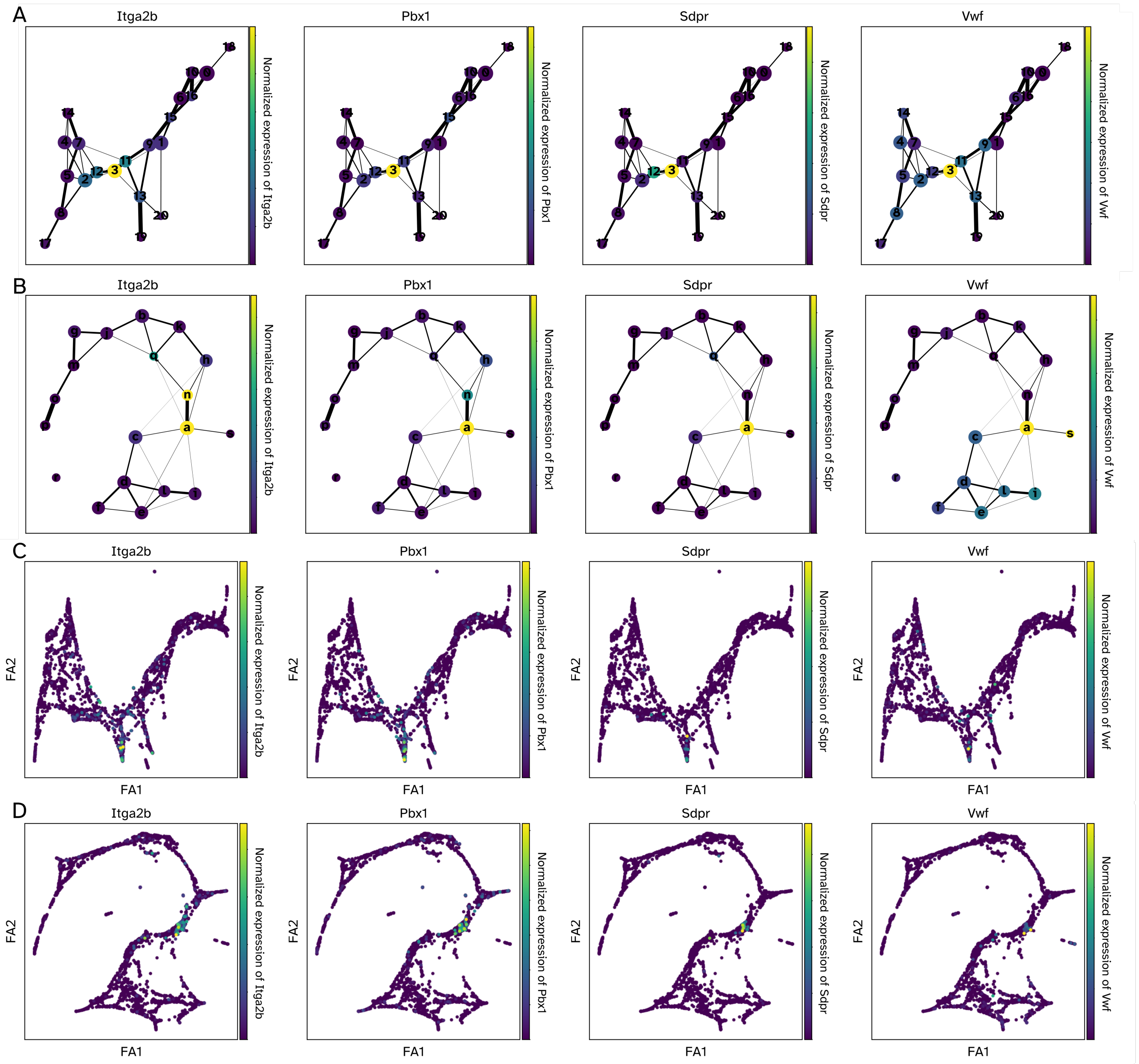


**Figure S12:** Megakaryocytes' marker genes activation patterns in experimental and simulated data. Panels A and C show the normalized gene expression of the marker genes in PAGA graphs for the simulated and the experimental data, respectively. Nodes correspond to Louvain clusters capturing discrete states and edges show transitions among these states. Panels B and D show PAGA-initialized single-cell embeddings obtained using ForceAtlas2. In all figures, colors yellow and purple show the highest and lowest normalized gene expression value, respectively.

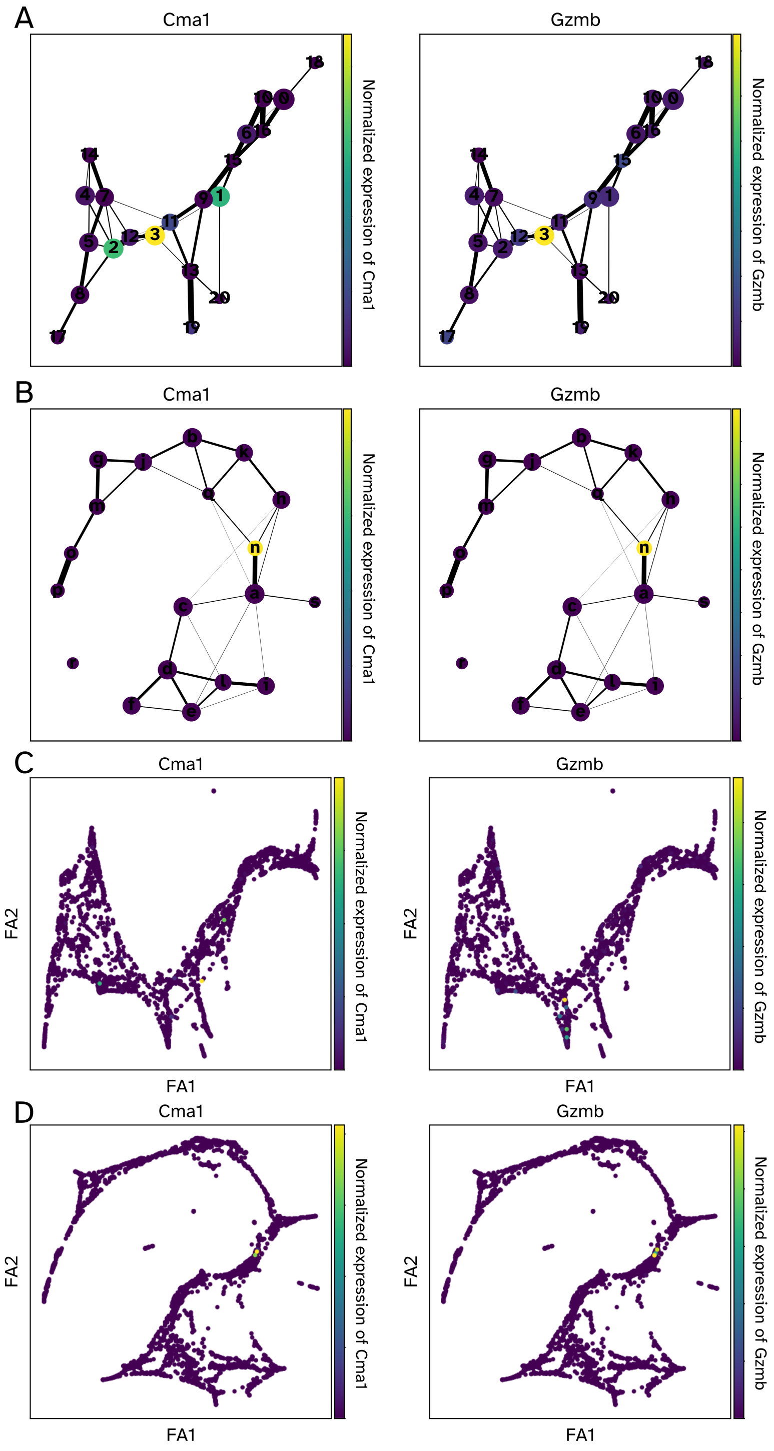


**Figure S13:** Mast cells' marker genes activation patterns in experimental and simulated data. Panels A and C show the normalized gene expression of the marker genes in PAGA graphs for the simulated and the experimental data, respectively. Nodes correspond to Louvain clusters capturing discrete states and edges show transitions among these states. Panels B and D show PAGA-initialized single-cell embeddings obtained using ForceAtlas2. In all figures, colors yellow and purple show the highest and lowest normalized gene expression value, respectively.

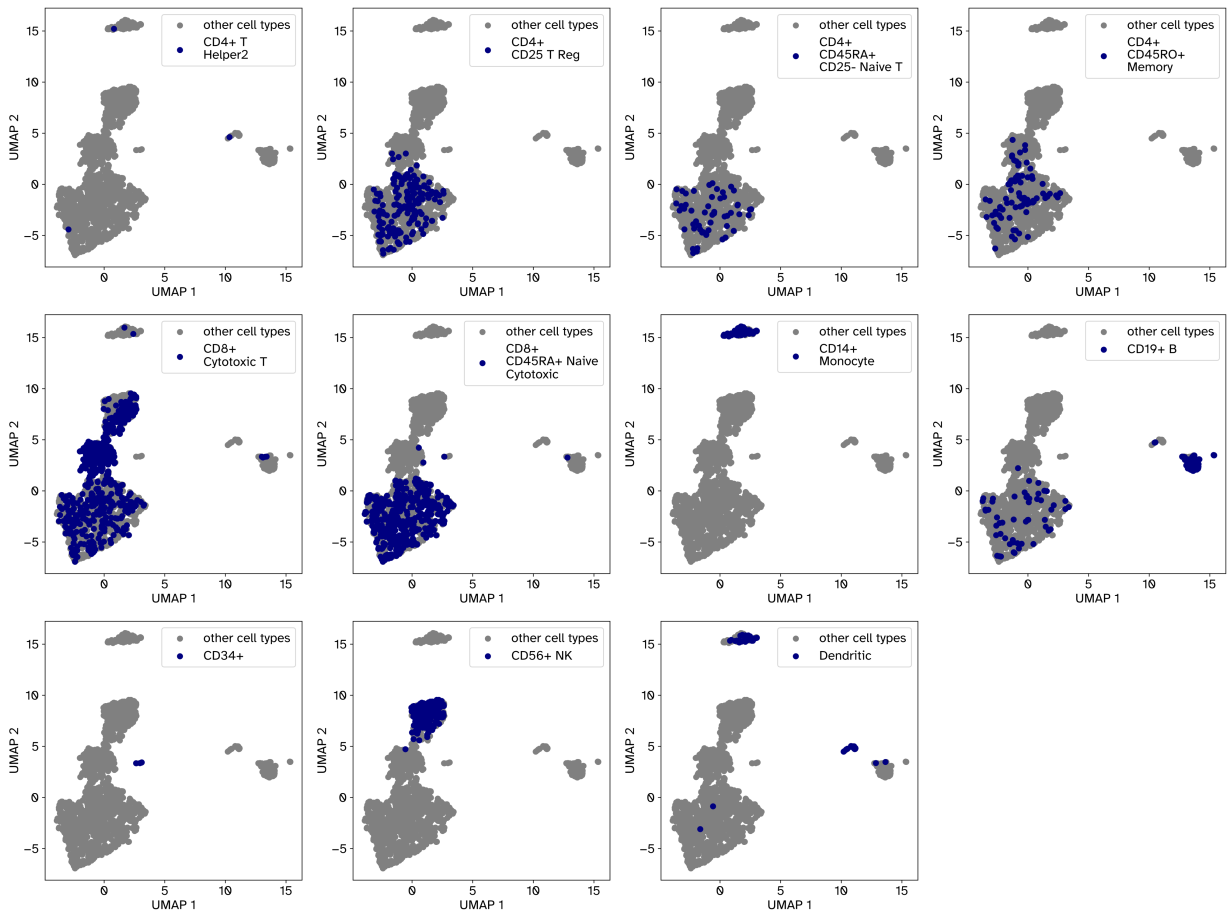


**Figure S14:** UMAP embedding of different cell types of the experimental PBMC-All dataset used for the analysis reported in Figure 6. We used cell-type annotations provided by 10x Genomics assigned by maximum correlation of PBMCs to filtered populations.

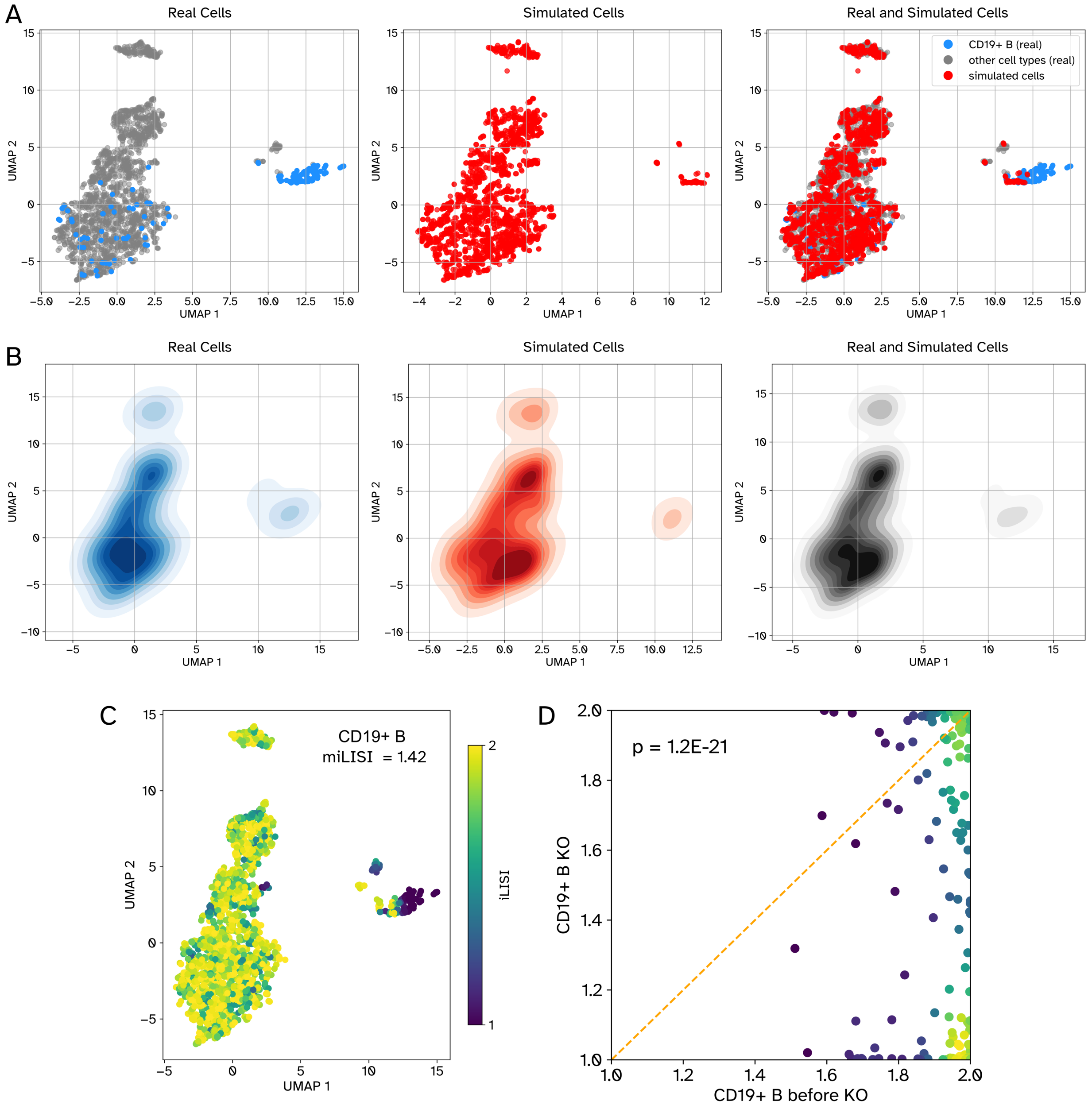


**Figure S15:** TF knockout analysis of CD 19+ B cells. Panels A and B show the UMAP and density plots of 2000 randomly selected cells from the experimental PBMC-All dataset (left), same number of simulated cells after knockout of top three TFs of CD19+ B cells (middle), and all together (right), respectively. TFs were omitted as features when generating UMAP plots. Panel C shows the iLISI value of each real cell (miLISI = 1.42) calculated from a UMAP embedding, jointly obtained from the experimental cells and the same number of simulated cells after knockout. D) The scatter plot shows the iLISI values of CD19+ B cells calculated along with unperturbed simulated cells (x-axis) and along with perturbed simulated cells (y-axis). The circles correspond to real cells and their colors reflect the density of datapoints in that region.

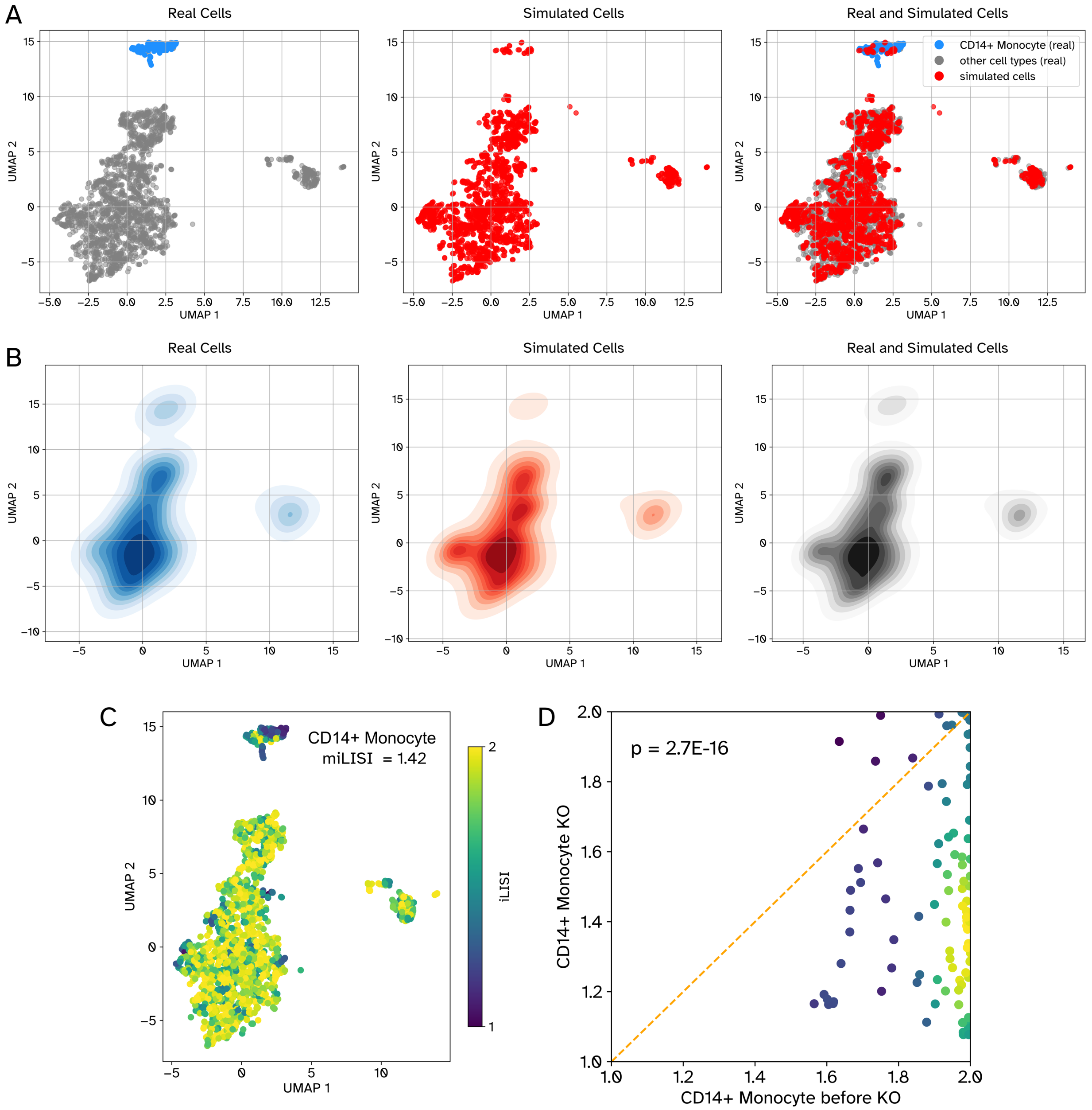


**Figure S16:** TF knockout analysis of CD14+ Monocyte cells. Panels A and B show the UMAP and density plots of 2000 randomly selected cells from the experimental PBMC-All dataset (left), same number of simulated cells after knockout of top three TFs of CD14+ Monocyte cells (middle), and all together (right), respectively. TFs were omitted as features when generating UMAP plots. Panel C shows the iLISI value of each real cell (miLISI = 1.42) calculated from a UMAP embedding, jointly obtained from the experimental cells and the same number of simulated cells after knockout. D) The scatter plot shows the iLISI values of CD14+ Monocyte cells calculated along with unperturbed simulated cells (x-axis) and along with perturbed simulated cells (y-axis). The circles correspond to real cells and their colors reflect the density of datapoints in that region.

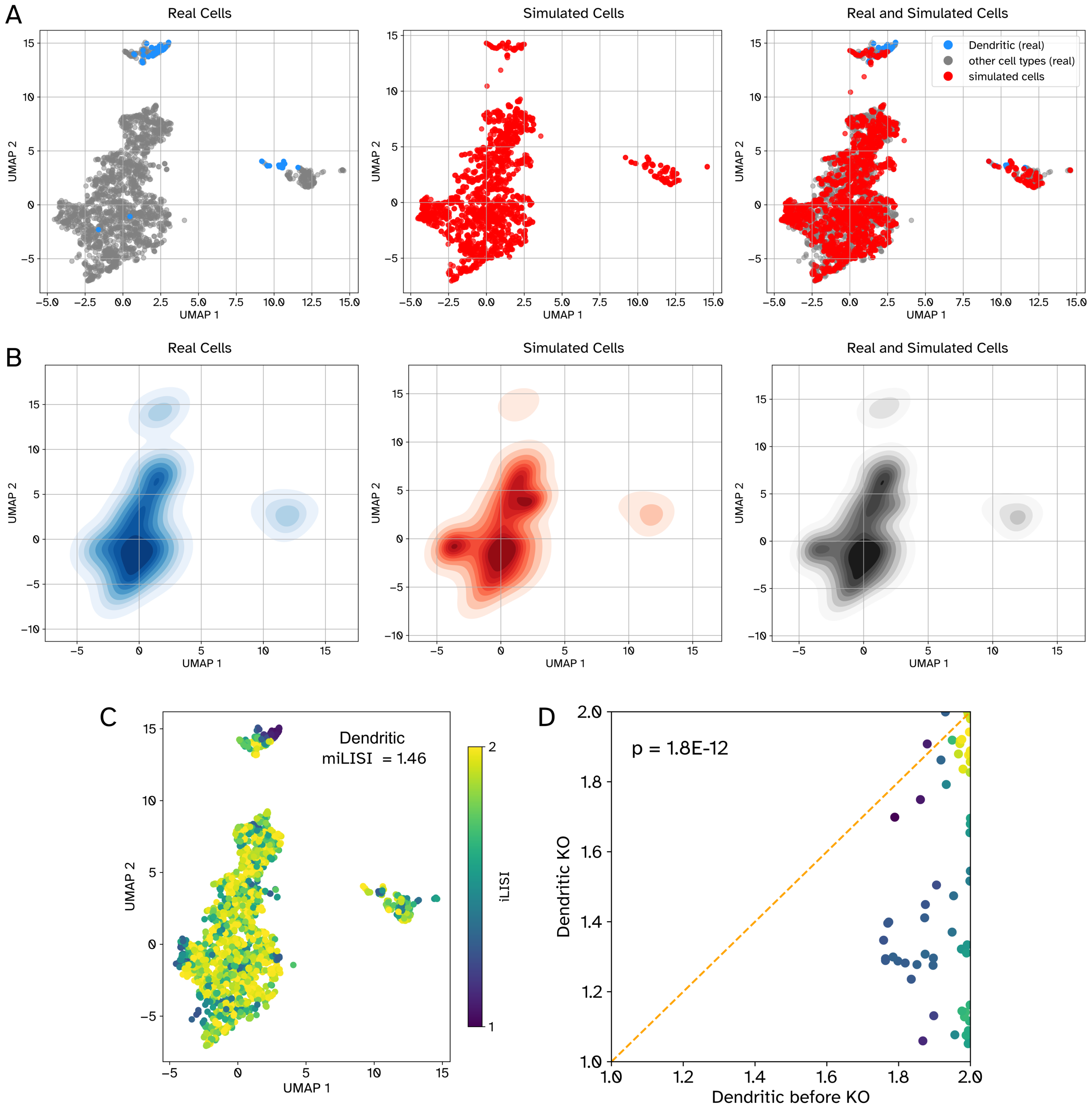


**Figure S17:** TF knockout analysis of Dendritic cells. Panels A and B show the UMAP and density plots of 2000 randomly selected cells from the experimental PBMC-All dataset (left), same number of simulated cells after knockout of top three TFs of Dendritic cells (middle), and all together (right), respectively. TFs were omitted as features when generating UMAP plots. Panel C shows the iLISI value of each real cell (miLISI = 1.46) calculated from a UMAP embedding, jointly obtained from the experimental cells and the same number of simulated cells after knockout. D) The scatter plot shows the iLISI values of Dendritic cells calculated along with unperturbed simulated cells (x-axis) and along with perturbed simulated cells (y-axis). The circles correspond to real cells and their colors reflect the density of datapoints in that region.

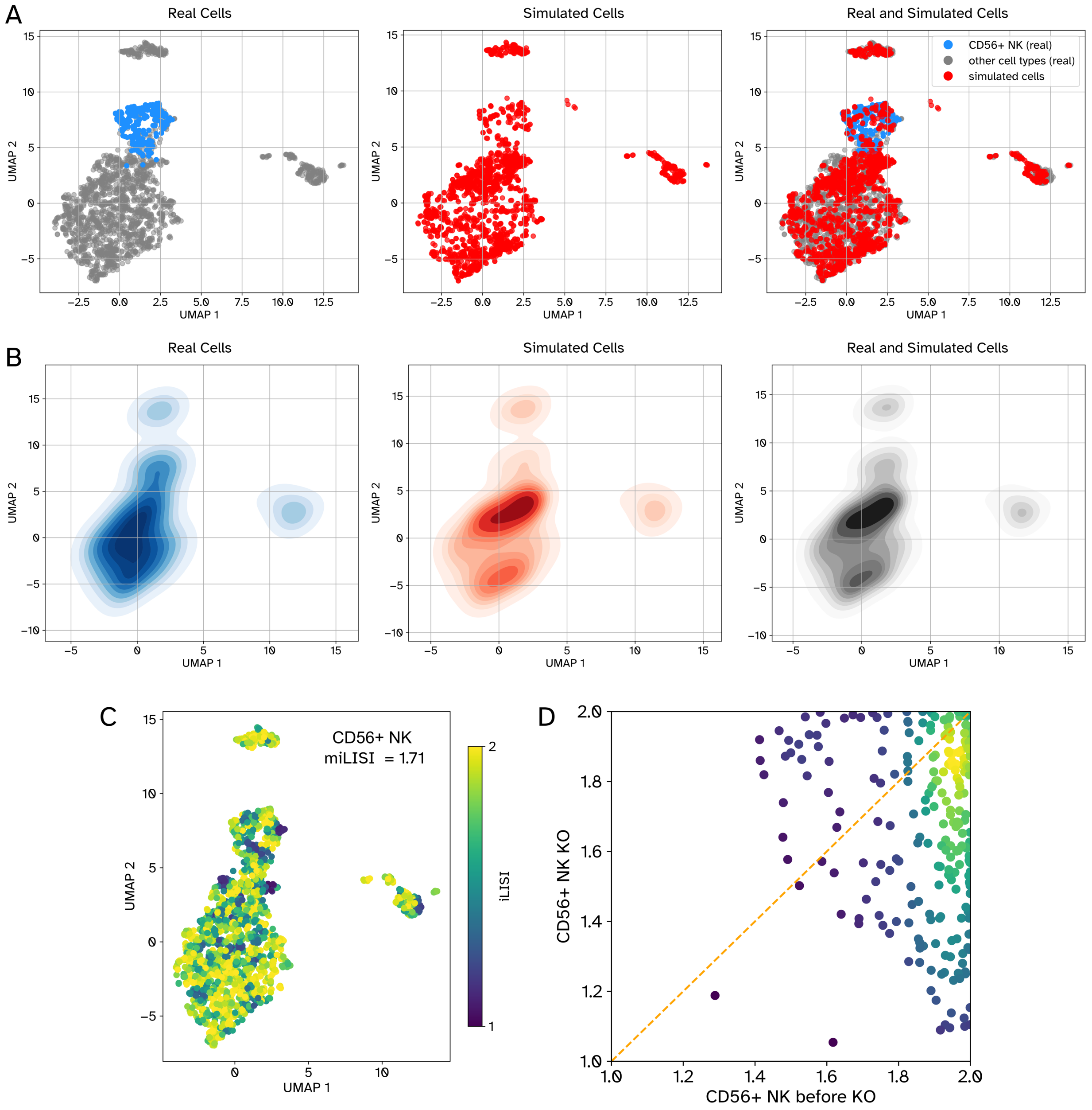


**Figure S18:** TF knockout analysis of CD56+ NK cells. Panels A and B show the UMAP and density plots of 2000 randomly selected cells from the experimental PBMC-All dataset (left), same number of simulated cells after knockout of top three TFs of CD56+ NK cells (middle), and all together (right), respectively. TFs were omitted as features when generating UMAP plots. Panel C shows the iLISI value of each real cell (miLISI = 1.71) calculated from a UMAP embedding, jointly obtained from the experimental cells and the same number of simulated cells after knockout. D) The scatter plot shows the iLISI values of CD56+ NK cells calculated along with unperturbed simulated cells (x-axis) and along with perturbed simulated cells (y-axis). The circles correspond to real cells and their colors reflect the density of datapoints in that region.

**References**


1 Che, T., Li, Y., Jacob, A. P., Bengio, Y. & Li, W. Mode regularized generative adversarial networks. *arXiv preprint arXiv:1612.02136* (2016).

2 Gulrajani, I., Ahmed, F., Arjovsky, M., Dumoulin, V. & Courville, A. C. Improved training of wasserstein gans. *Advances in neural information processing systems* **30** (2017).

3 Arjovsky, M. & Bottou, L. Towards principled methods for training generative adversarial networks. *arXiv preprint arXiv:1701.04862* (2017).

4 Kushwaha, V. & Nandi, G. in *2020 IEEE 4th Conference on Information & Communication Technology (CICT).* 1-6 (IEEE).

5 Liu, K., Tang, W., Zhou, F. & Qiu, G. in *Proceedings of the IEEE/CVF international conference on computer vision.* 6382-6390.

6 Yao, Y., Pan, Y., Tsang, I. W. & Yao, X. in *International Conference on Neural Information Processing.* 40-48 (Springer).

7 Arjovsky, M., Chintala, S. & Bottou, L. in *International conference on machine learning.* 214-223 (PMLR).

8 Villani, C. *Optimal transport: old and new*. Vol. 338 (Springer, 2009).

9 Wolf, F. A. *et al.* PAGA: graph abstraction reconciles clustering with trajectory inference through a topology preserving map of single cells. *Genome biology* **20**, 1-9 (2019).
